# Early Polycomb-target deregulations in Hutchinson-Gilford Progeria Syndrome revealed by heterochromatin analysis

**DOI:** 10.1101/799668

**Authors:** Endre Sebestyén, Fabrizia Marullo, Federica Lucini, Andrea Bianchi, Cristiano Petrini, Sara Valsoni, Ilaria Olivieri, Laura Antonelli, Francesco Gregoretti, Gennaro Oliva, Francesco Ferrari, Chiara Lanzuolo

**Affiliations:** IFOM, the FIRC Institute of Molecular Oncology, Milan, Italy; Institute of Cell Biology and Neurobiology, National Research Council, Rome, Italy; Istituto Nazionale Genetica Molecolare “Romeo ed Enrica Invernizzi”, Milan, Italy; IRCCS Santa Lucia Foundation, Rome, Italy; Institute for High Performance Computing and Networking, Naples, Italy; Institute of Molecular Genetics, National Research Council, Pavia, Italy; Institute of Biomedical Technologies, National Research Council, Milan, Italy

**Keywords:** Heterochromatin, Genome structure, Lamin A/C, Progeria, Lamin Associated Domains, Transcription regulation

## Abstract

Hutchinson-Gilford progeria syndrome (HGPS) is characterized by the progressive accumulation of progerin, an aberrant form of Lamin A. This leads to chromatin structure disruption, in particular by interfering with Lamina Associated Domains. Although several cellular and molecular alterations have been characterized, it is still unclear how chromatin structural changes translate into premature senescence in HGPS. Moreover, early events in chromatin remodeling have not been detected so far. We developed a new high-throughput sequencing-based method, named SAMMY-seq, for genome-wide characterization of heterochromatin accessibility changes. Using SAMMY-seq, we detected early stage alterations of chromatin structure in HGPS primary fibroblasts. Of note, these structural changes do not disrupt the distribution of H3K9me3 but are associated with site-specific H3K27me3 variations and transcriptional dysregulation of Polycomb target genes. Our results show that SAMMY-seq represents a novel and sensitive tool to characterize heterochromatin alterations. Moreover, we found that the assembly of lamin associated domains is strictly connected to the correct Polycomb repression, rapidly lost in early HGPS pathogenesis.

---

In eukaryotic nuclei, DNA shows different levels of compaction recognizable by electron microscopy, known as euchromatin and heterochromatin (reviewed in^1,2^). The more accessible and less condensed euchromatin is generally enriched in expressed genes. Whereas heterochromatin contains highly condensed DNA, including pericentromeric and telomeric regions^3–5^, and genomic regions with unique packaging properties targeted by the Polycomb-group proteins (PcG)^6^. These proteins are developmentally regulated transcriptional factors acting as parts of multimeric complexes such as Polycomb Repressive Complex 1 and 2 (PRC1 and PRC2)^7^.

Another key player of chromatin organization is the nuclear lamina (NL). NL is mainly composed of A- and B-type lamins, intermediate filaments of 3.5 nm thickness which form a dense network at the nuclear periphery^8,9^. NL preferentially interacts with the genome at specific regions called Lamina Associated Domains (LADs), with sizes ranging from 100Kb to 10Mb^10^. LADs are enriched in H3K9me2 and H3K9me3 histone modifications that are typical of inactive heterochromatic regions. LAD borders are marked by the PRC2-dependent H3K27me3 histone mark^11–13^, which is characteristic of inactive PcG-regulated chromatin regions. The ensemble of lamins, chromatin marks and PcG factors around LADs creates a repressive environment^14,15^, with heterochromatin and PcG target regions adjacent to each other^16,17^. In line with these observations, we previously demonstrated that Lamin A functionally interacts with PcG^18,19^, as later also reported by others^17,20–22^.

In physiological conditions, the association or detachment of LADs from the NL are related to the spatiotemporal regulation of gene expression in cell differentiation processes^13,23,24^. The crucial function of these dynamics is attested by an entire class of genetic diseases, called laminopathies, where specific components of the NL are altered^13,21,25^. Among them, Hutchinson-Gilford Progeria Syndrome (HGPS, OMIM 176670) is a rare and fatal disease caused by a *de novo* point mutation in the LMNA gene^26^, resulting in a truncated splicing mutant form of Lamin A protein, known as progerin. The disease phenotype is characterized by early onset of several symptoms resulting in a severe premature aging in pediatric patients. At the cellular level, the accumulation of progerin induces progressive alterations of nuclear shape and cellular senescence^27^. Notably, inhibition of PcG functions can trigger “nuclear blebbing”, an alteration of the nuclear shape typically observed in Lamin A deficient cells^28^, suggesting that chromatin architecture and nuclear shape strictly depend on each other. Consistently, LADs organization and PcG-dependent H3K27me3 mark are altered in HGPS^29,30^. However, while extensive disruption of chromatin structure has been reported by Hi-C in late-passage HGPS fibroblasts, this could not be detected in early passage cells, commonly considered a model to identify early molecular alterations in the disease^29^.

In recent years, the widespread adoption of experimental techniques based on high-throughput sequencing (NGS) has been instrumental in advancing the knowledge of chromatin structure and function^31^ ^32^. Notably, several of these techniques were originally designed to map active chromatin regions, including ChIP-seq^33^, ATAC-seq^34^, DNase-seq^35^ and MNase-seq^36^. On the other hand, few options are available for genome-wide mapping of heterochromatin and LADs. These include ChIP-seq or DamID-seq^37^ targeting NL components or ChIP-seq for the heterochromatin associated histone marks (H3K9 methylation). Such techniques suffer major limitations as ChIP-seq relies on crosslinking, causing technical biases, and antibodies, with several potential issues of sensitivity or cross-reactivity. Furthermore, DamID-seq relies on the exogenous expression of a transgene and it can’t be applied on primary cells or tissues.

Here, we present a novel high-throughput sequencing based method to map lamina associated heterochromatin regions. This technique relies on the sequential isolation and sequencing of multiple chromatin fractions, enriched for differences in accessibility. Our method is robust, fast, easy-to-perform, it does not rely on crosslinking or antibodies, and it can be applied on primary cells. In normal human fibroblasts, we can reliably identify genomic regions with heterochromatin marks (H3K9me3) and LADs.

In early-passage HGPS fibroblasts we found patient-specific changes of chromatin accessibility. Although HGPS chromatin remodeling is not accompanied by H3K9me3 changes, differentially compacted domains affect H3K27me3 distribution and the transcriptional state of PcG regulated bivalent genes. These findings suggest that chromatin structural changes are an early event in HGPS nuclear remodeling and interfere with proper PcG control. We named our technology SAMMY-seq (Sequential Analysis of MacroMolecules accessibilitY), as a tribute to Sammy Basso, the founder of the Italian Progeria Association, a passionate and inspiring advocate of research on laminopathies.

## Results

### SAMMY-seq allows mapping of lamina associated heterochromatic regions

We developed the SAMMY-seq method for genome-wide mapping of chromatin fractions separated by accessibility. The method is based on the sequential extraction of distinct nuclear fractions^18,38,39^ containing: soluble proteins (S1 fraction), DNase-sensitive chromatin (S2 fraction), DNase-resistant chromatin (S3 fraction) and the most condensed and insoluble portion of chromatin (S4 fraction) (Fig. 1a and Supplementary Fig. 1a). We further adapted the method to leverage high throughput DNA sequencing for genome-wide mapping of the distinct chromatin fractions (see Methods).

**Figure 1.**
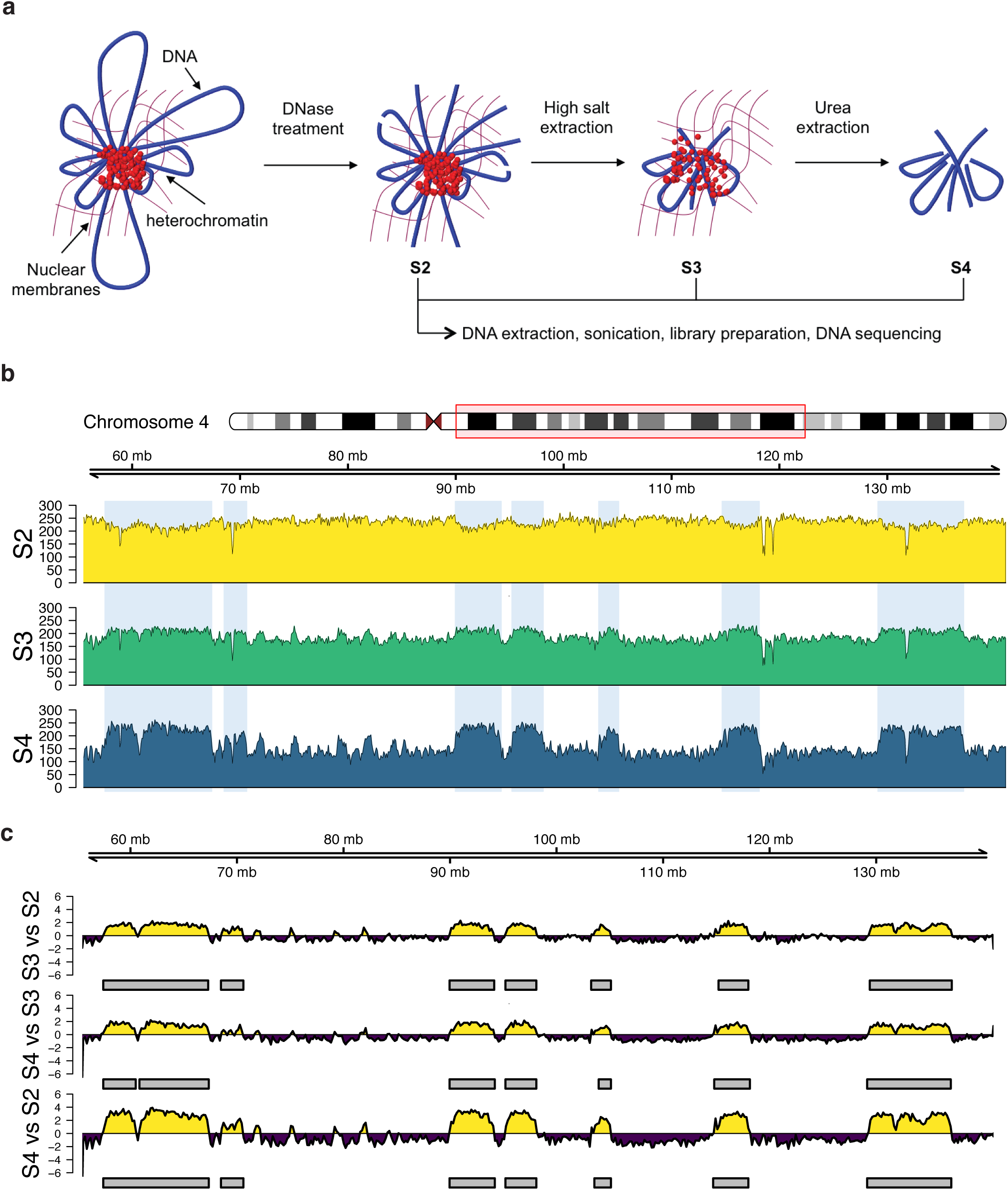
SAMMY-seq isolates specific DNA regions in fibroblast samples. **a**, Schematic representation of SAMMY-seq. Chromatin fractions are sequentially isolated after DNase treatment, high salt and urea buffers extractions. The associated genomic DNA is extracted, sonicated and processed for high-throughput sequencing. **b**, Distribution of SAMMY-seq reads along a representative region (85Mb region in chr4:55480346-141005766). Library size normalized read counts over 10Kb genomic bins are shown for each sequenced fraction of a control fibroblast sample (CTRL002). **c**, Differential reads distribution across pairwise comparisons of SAMMY-seq fractions in a control fibroblast sample (CTRL002) along the genomic region in (a). Less accessible fractions are compared to more accessible ones used as reference (S3 vs S2; S4 vs S3; S4 vs S2). Regions of signal enrichment or depletion over the reference samples are marked in yellow or purple, respectively. The smoothed differential signal is calculated with SPP, and significantly enriched regions (SAMMY-seq domains) are called with EDD (see Online methods for details) and reported as grey boxes under the enrichment signal track.

We first applied SAMMY-seq to 3 independent normal skin primary fibroblast cell lines, originating from 3 different individuals (control samples). The average number of reads in the different fractions was 72 million (S2), 62 million (S3) and 65 (S4) million (Fig. 1b, Supplementary Fig. 1b, Supplementary Table 1), and on average 58% (S2), 54% (S3) and 52% (S4) of the genome was covered by at least one read (Supplementary Fig. 1c). The genome-wide correlation of read coverage confirms a high reproducibility of signal for the S3 and S4 fractions, enriched for more compacted chromatin, whereas the S2 fraction is more variable across biological replicates (Supplementary Fig. 1d). We noticed that correlation computed over genomic bins of variable sizes, ranging from 10Kb to 5Mb, becomes more stable with larger bin sizes (Supplementary Fig. 1e). At 1Mb resolution, the mean Spearman correlation is 0.27 for the S2 fraction, 0.92 for S3, and 0.79 for S4. Visual inspection of read coverage profiles shows megabase-scale “bumps” in S3 and, more prominently, in S4 fractions (Fig. 1b). To detect relative enrichment of specific genomic regions between chromatin fractions, we applied the Enriched Domain Detector (EDD)^40^ algorithm to our data. EDD was specifically designed to identify broad regions of enrichment in high-throughput sequencing results. We looked for differentially enriched genomic regions using pairwise comparisons of less accessible vs more accessible fractions used as baseline: S3 vs S2, S4 vs S3 and S4 vs S2 (Fig. 1c). We found a well-defined set of regions (SAMMY-seq domains) consistently enriched in the less accessible fractions. These domains are conserved across samples, with the S4 vs S2 comparison being the most consistent, and showing an average of 70.18% conservation (Supplementary Fig. 1f). To confirm the robustness of our results we repeated the differential enrichment analysis after down-sampling to 50% of sequencing reads. We obtained similar SAMMY-seq domains, with an average overlap of 80.64 % for S3 vs S2, 82.22 % for S4 vs S3 and 83.42 % for S4 vs S2 across samples (Supplementary Fig 1g). Further down-sampling to 25% of reads still led to an average of 61.92 % overlapping domains across samples and comparisons (Supplementary Fig 1g). SAMMY-seq domains have an average size of 2,26 Mb and cover on average 527 Mb (Supplementary Fig. 1h, i, Supplementary Table 2). Of note, other chromatin fractionation-based NGS methods have been previously described^41,42^, however they are not directly comparable to our technology as they adopt different conditions, enzymes and cellular models.

To functionally characterize the SAMMY-seq enriched regions, we compared the genome-wide read coverage with reference chromatin mark profiles for the same cell type. Using data from the Roadmap Epigenomics consortium^43^ and other publications^29,44,45^ we noticed that SAMMY-seq signal (S4 vs S2 enrichment) is highly consistent across biological replicates (CTRL002, CTRL004, CTRL013), it is inversely correlated with open chromatin marks (ATAC-seq and DNase-seq) and is positively correlated with Lamin A/C and B ChIP-seq signal (Fig. 2a). We confirmed this is a consistent genome-wide pattern using StereoGene kernel correlation^46^ for the unbiased comparison of different types of chromatin marks (Fig. 2b). Similarly, when considering histone modifications, we observed a positive association with heterochromatin mark (H3K9me3) and inverse correlation with active chromatin (H3K4me1) as well as Polycomb-regulated chromatin (H3K27me3) (Fig. 2a-b). The latter observation is compatible with previous reports that H3K27me3 is located at the border of heterochromatic LAD regions^11–13,17^. The genome-wide correlation patterns were also consistent when considering individual SAMMY-seq fractions (Supplementary Fig. 2a), confirming that we were progressively enriching for more compact chromatin in the sequential extraction steps.

**Figure 2.**
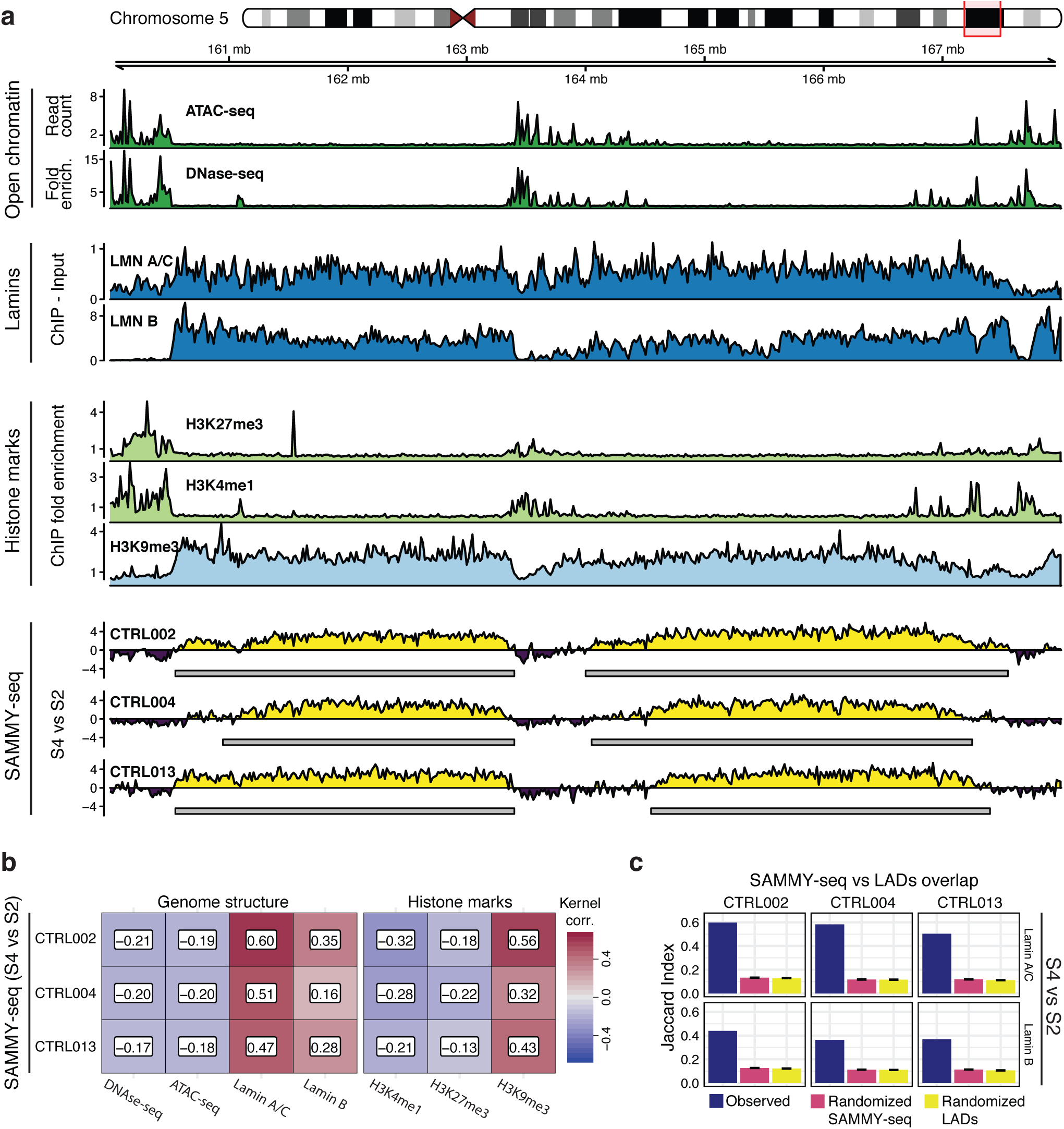
SAMMY-seq preferentially isolates heterochromatic domains. **a**, Visualization of SAMMY-seq enrichment signal along with multiple chromatin marks on a representative region (8Mb region in chr5:158000000-170000000). From top to bottom: tracks for open chromatin (ATAC-seq and DNase-seq – dark green); association to lamina (Lamin A/C and Lamin B ChIP-seq – dark blue); ChIP-seq for histone marks associated to PcG regulation (H3K27me3 – light green), active chromatin (H3K4me1 – light green) or heterochromatin (H3K9me3 – light blue); SAMMY-seq enrichment signal (S4 vs S2) in 3 control samples (yellow or purple colored track for enrichment or depletion, respectively, over the S2 reference baseline). Grey boxes under each SAMMY-seq track show the SAMMY-seq domains. See Online methods for details of each type of signal processing. **b**, Genome-wide kernel correlation (StereoGene) for SAMMY-seq of all control samples (S4 vs S2 enrichment) compared to ATAC-seq, DNase-seq, and ChIP-seq (Lamin A/C, Lamin B, H3K27me3, H3K4me1 and H3K9me3). **c**, Overlap (Jaccard Index - JI) of control samples SAMMY-seq domains (S4 vs S2) with LADs (Lamin A/C or Lamin B ChIP-seq enrichment domains). The observed JI (blue bar) is compared to mean JI across 10,000 randomizations of SAMMY-seq domains (pink bar) or LADs (yellow bar) positions along the genome. All Jaccard Indexes are significantly higher than the randomized values (all empirical p-values < 0.0001). The black rectangle over the randomized values shows the +/− 2 standard error range.

To quantify the overlap between SAMMY-seq enrichment regions and LADs, we used Lamin ChIP-seq datasets (Lamin A/C and Lamin B)^29,44^ and compared them with SAMMY-seq domains (S4 vs S2, S4 vs S3, S3 vs S2). On average 79% of SAMMY-seq domains across different comparisons and samples overlap with Lamin A/C LADs (Supplementary Table 3). All of the comparisons showed a significant overlap (average Jaccard Index 0.56 for Lamin A/C in S4 vs S2), as assessed by randomizing either LADs or SAMMY-seq domains (Fig. 2c and Supplementary Fig. 2b). Interestingly, the overlap with Lamin A/C associated domains was 21% higher than with Lamin B associated domains, suggesting that SAMMY-seq preferentially enriches for regions interacting with Lamin A/C. This is also consistent with the observed higher genome-wide correlation with Lamin A/C compared to Lamin B (Fig. 2b and Supplementary Fig. 2a). As a further confirmation that SAMMY-seq specifically detects the most compact heterochromatin, such as the lamina associated heterochromatic regions, we observed that SAMMY-seq domains have an even higher overlap with regions defined as both LADs and H3K9me3 enriched (77.84% average overlap), rather than considering each of the two markers independently (64.9% and 42.69% average overlap, respectively) (Supplementary Fig. 2c).

### SAMMY-seq detects chromatin changes in early passage progeria fibroblasts

We applied SAMMY-seq on early passage HGPS fibroblasts, a cellular model to identify early molecular alterations in Progeria, as progerin accumulates during passaging (Supplementary Fig. 3a). According to recent literature, primary HGPS fibroblasts entering senescence show alterations in DNA interactions with Lamin A/C^29^. However, at early passages Hi-C experiments for genome-wide mapping of 3D chromatin architecture failed to detect alterations in the pairwise contact frequency between genomic loci. These were evident only at later-passage cells, representing more advanced stages of senescence, when cells exhibit severe nuclear blebs and inhibited proliferation^29^.

We confirmed that early-passage HGPS fibroblasts do not show senescence-associated features such as beta-galactosidase positivity (Supplementary Fig. 3b), reduction in the proliferation rate (Supplementary Fig. 3c), changes in nuclear area and morphology (Supplementary Fig. 3d, e, f). We used SAMMY-seq to investigate chromatin changes in early-passage (passage number from 10 to 12) skin fibroblasts from 3 independent HGPS patients (Supplementary Fig. 1a, b). Compared with analysis showing a consistent overlap (S4 vs S2) across control replicates, in HGPS patients the SAMMY-seq domains appeared scattered and with generally lower signal-to-noise ratio (Fig. 3a). This would be compatible with a general loss of chromatin organization, resulting in more random distribution of genomic regions among the different fractions. Pairwise comparisons confirmed that SAMMY-seq domains are largely overlapping among control samples (average Jaccard Index 0.67) whereas this overlap is lost in progeria samples, most notably in the S4 vs S2 comparison (average Jaccard Index 0.11) (Fig. 3b and Supplementary Table 4). Moreover, SAMMY-seq enrichment in control samples is matching lamins and H3K9me3 ChIP-seq enrichment, whereas this pattern is lost in HGPS SAMMY-seq samples (Fig. 3a). We confirmed this is a genome-wide trend by analyzing chromatin marks at SAMMY-seq domain borders (Fig. 3c). SAMMY-seq domains from control samples (S4 vs S2, S4 vs S3, S3 vs S2) correspond to regions normally enriched in Lamin A/C, Lamin B and H3K9me3 ChIP-seq reads, as opposed to flanking regions outside of domain borders. Progeria SAMMY-seq domains (S3 vs S2) are similar to controls. However, when considering more compacted S4 fraction, the association to reference LADs and heterochromatin marks is lost in 2 patients for the S4 vs S2 and 3 patients for the S4 vs S3 comparisons, respectively (Fig. 3c). Here, both the lamins and the H3K9me3 signals are flattened out suggesting no specific pattern.

**Figure 3.**
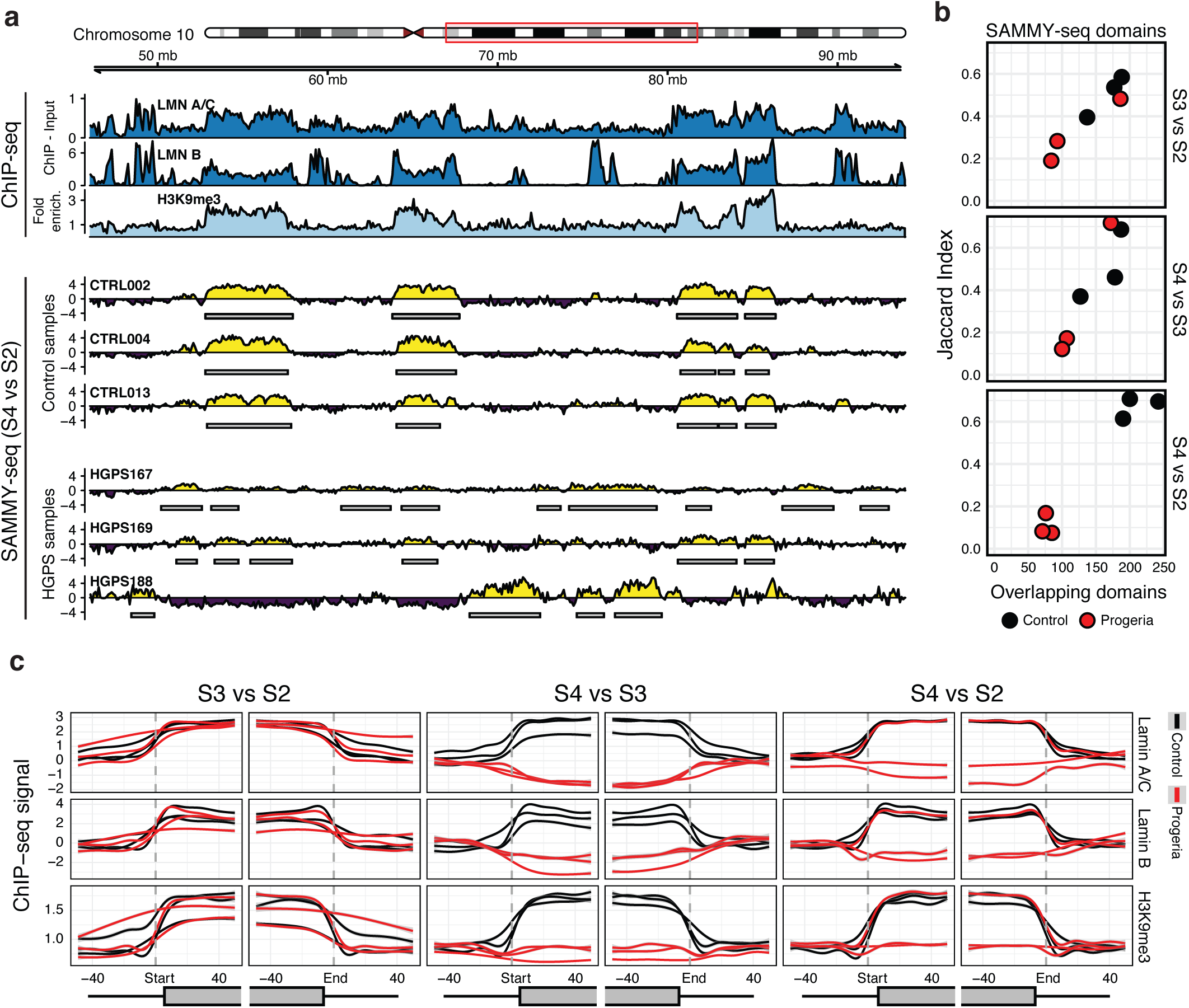
HGPS fibroblasts show early changes in chromatin accessibility. **a**, Genomic tracks for SAMMY-seq in 3 control and 3 HGPS samples along with chromatin marks in a representative region (48 Mb region in chr10:46000000-94000000). From top to bottom: tracks for association to lamina (Lamin A/C and Lamin B ChIP-seq – dark blue); heterochromatin (H3K9me3 ChIP-seq – light blue); SAMMY-seq enrichment signal (S4 vs S2) in 3 control and 3 HGPS samples (yellow or purple colored track for enrichment or depletion, respectively, over the S2 reference baseline). Grey boxes under each SAMMY-seq track show the SAMMY-seq domains. **b**, Pairwise overlap of SAMMY-seq domains (S3 vs S2, S4 vs S3, S4 vs S2) between control (black dots) or HGPS (red dots) sample pairs (JI on y-axis, number of overlapping regions on x-axis). Upper-tail Fisher-test p-value < 0.01 for all overlaps, except for HGPS S4 vs S2 pairwise overlaps. **c**, Smoothed average ChIP-seq enrichment signal for chromatin marks around SAMMY-seq domain borders. Average ChIP-seq signal for Lamin A/C (upper row), Lamin B (middle row) and H3K9me3 (bottom row) is reported around the SAMMY-seq domain borders start (left side plots) or end (right side plots) for domains detected in each control (black lines) and progeria (red lines) sample. Results for each set of SAMMY-seq enrichment domains are reported (S3 vs S2 on the left, S4 vs S3 center, S4 vs S2 on the right) using a +/-50 bins window (10Kb bin size) centered on the start or end domain border positions (vertical dashed grey line).

### Early changes of chromatin structure in HGPS fibroblasts is not accompanied by alterations of H3K9me3 patterns

To further investigate how changes in chromatin accessibility in early-passage HGPS fibroblasts affect the epigenetic regulation, we examined H3K9me3 distribution by ChIP-seq on the same set of control and HGPS fibroblasts analyzed by SAMMY-seq (Fig. 4a). Visual exploration of sequencing reads distribution revealed that, despite chromatin remodeling was detected by SAMMY-seq, H3K9me3 profiles were unaffected in HGPS. H3K9me3 enriched regions are largely conserved between controls and HGPS cells as shown by the genome-wide overlap (between 69% and 80%) (Fig. 4b). Analysis of H3K9me3 patterns at SAMMY-seq domain borders (Supplementary Fig. 4a) and Jaccard index score (Fig. 4c and Supplementary Fig. 4b) confirmed that only controls exhibit a correlation between SAMMY-seq and H3K9me3.

**Figure 4.**
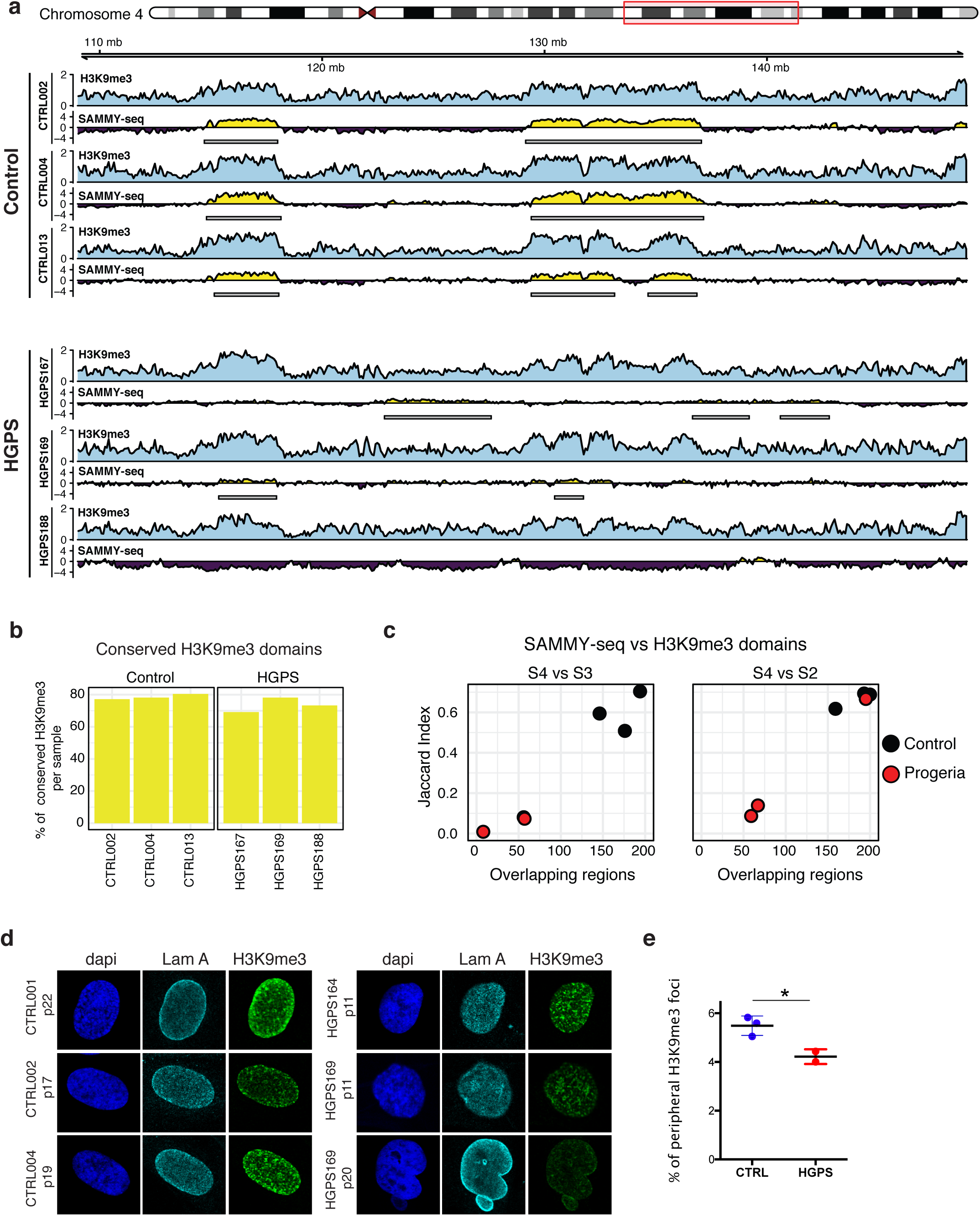
Early chromatin structure disruption in progeria is not accompanied by alterations in H3K9me3. **a**, Genomic tracks for paired H3K9me3 ChIP-seq and SAMMY-seq in 3 control and 3 HGPS samples for a representative region (40Mb region in chr4:109000000-149000000). Enrichment signals (computed using SPP) for each sample are shown (H3K9me3 ChIP-seq – light blue; yellow or purple colored track for SAMMY-seq S4 vs S2 enrichment or depletion, respectively). Grey boxes under each SAMMY-seq track show the SAMMY-seq domains. **b**, Percentage of H3K9me3 domains conserved across all 3 control or all 3 progeria samples (computed as percent over per sample total H3K9me3 domains size). **c**, Overlap of H3K9me3 enriched domains and SAMMY-seq domains (S4 vs S3, S4 vs S2) for control (black dots) or HGPS (red dots) samples (JI on y-axis, number of overlapping regions on x-axis). All overlaps are significant in controls (for both S4 vs S3 and S4 vs S2 SAMMY-seq domains) (upper-tail Fisher-test p-value < 0.01 for all overlaps) and in HGPS169 (only for S4 vs S2 SAMMY-seq domains). **d**, Representative images of H3K9me3/Lamin A immunofluorescence analysis on control or HGPS fibroblasts. **e**, Percentage of H3K9me3 foci localized within 1 um from nuclear periphery in distinct cell populations. Comparisons were done using a two-tailed t-test. Statistically significant differences are marked * p < 0.05

In line with genomic data, at cellular level early-passage HGPS cells and controls showed comparable H3K9me3 global levels measured by western blot (Supplementary Fig. 4c, d). Immunofluorescence analysis with H3K9me3 antibody revealed that heterochromatin foci were detected in both controls and early-passage HGPS fibroblasts with high variability across samples (Fig. 4d) but without significant differences in their number per cell (Supplementary Fig. 4e). Instead, their intranuclear localization was different, with a shift towards the nuclear center in HGPS (Fig. 4d, e), supporting the hypothesis of heterochromatin detachment from the nuclear periphery (Fig. 3). Notably, as previously shown^30^, later passages (p20) HGPS fibroblasts showed an aberrant pattern of H3K9me3 foci (Fig. 4d), further indicating that chromatin remodeling in HGPS precedes H3K9me3 loss.

### Large-scale chromatin structural changes do not affect expression of LAD-associated genes

Using RNA-seq on the same set of control and HGPS fibroblasts we assessed the expression at the gene or transcript level detecting respectively 260 differentially expressed genes and 256 transcripts, suggesting that early stages of senescence in HGPS cells do not cause large scale changes in the transcriptome (Supplementary Fig. 5a). To clarify whether chromatin structural changes detected with SAMMY-seq have an effect on transcription, we examined the expression of protein coding genes located in SAMMY-seq domains. As expected^16^, genes within SAMMY-seq domains (S4 vs S2) have lower expression levels than those outside in all control samples (Supplementary Fig. 5b). More specifically, in the genomic regions flanking SAMMY-seq domain borders we noticed a clear pattern with a transition from lower expression within the domain to higher expression outside in all of the control samples (Supplementary Fig. 5c). A similar pattern was observed in HGPS samples when considering the same genomic regions (Supplementary Fig. 5b, c). This suggests that heterochromatin detachment from NL seen in HGPS (Fig 3 and 4e) was not sufficient to trigger gene expression deregulation inside SAMMY-seq domains, where H3K9me3 is also largely unchanged as shown above (Fig. 4a, b). Further confirming this hypothesis, the transcriptional activity within H3K9me3 domains of each sample is repressed in HGPS as expected (Supplementary Fig. 5d).

### HGPS chromatin remodeling affects expression of PcG regulated bivalent genes

As we noticed that each HGPS sample has unique SAMMY-seq domains (Fig.3), we compared individual HGPS expression profiles against the group of three controls to account for sample-specific patterns (see Online methods). To verify which biological functions are affected we performed a pathway enrichment analysis using the MSigDB database^47^ and selected the pathways commonly deregulated in all HGPS samples (Fig. 5a). Many of them are related to stem cells and cancer, in line with the hypothesis that accelerated aging affect stem cell niches^48^. Interestingly, we also found several pathways related to PcG regulation (H3K27me3 and PRC2) (Fig. 5a). To further investigate PcG role in early passage HGPS fibroblasts, we examined the total amounts of PRC1-subunit Bmi1 and PRC2-subunit Ezh2 (Supplementary Fig. 6a, b), as well as H3K27me3 levels (Supplementary Fig. 6c, d), which showed no significant differences, despite some level of inter-individual variability. We also checked intranuclear architecture of Ezh2 (Supplementary Fig. 6e, f) and H3K27me3 foci (Supplementary Fig. 6g, h) and we found that their number and size do not change at early-passage HGPS cells.

**Figure 5.**
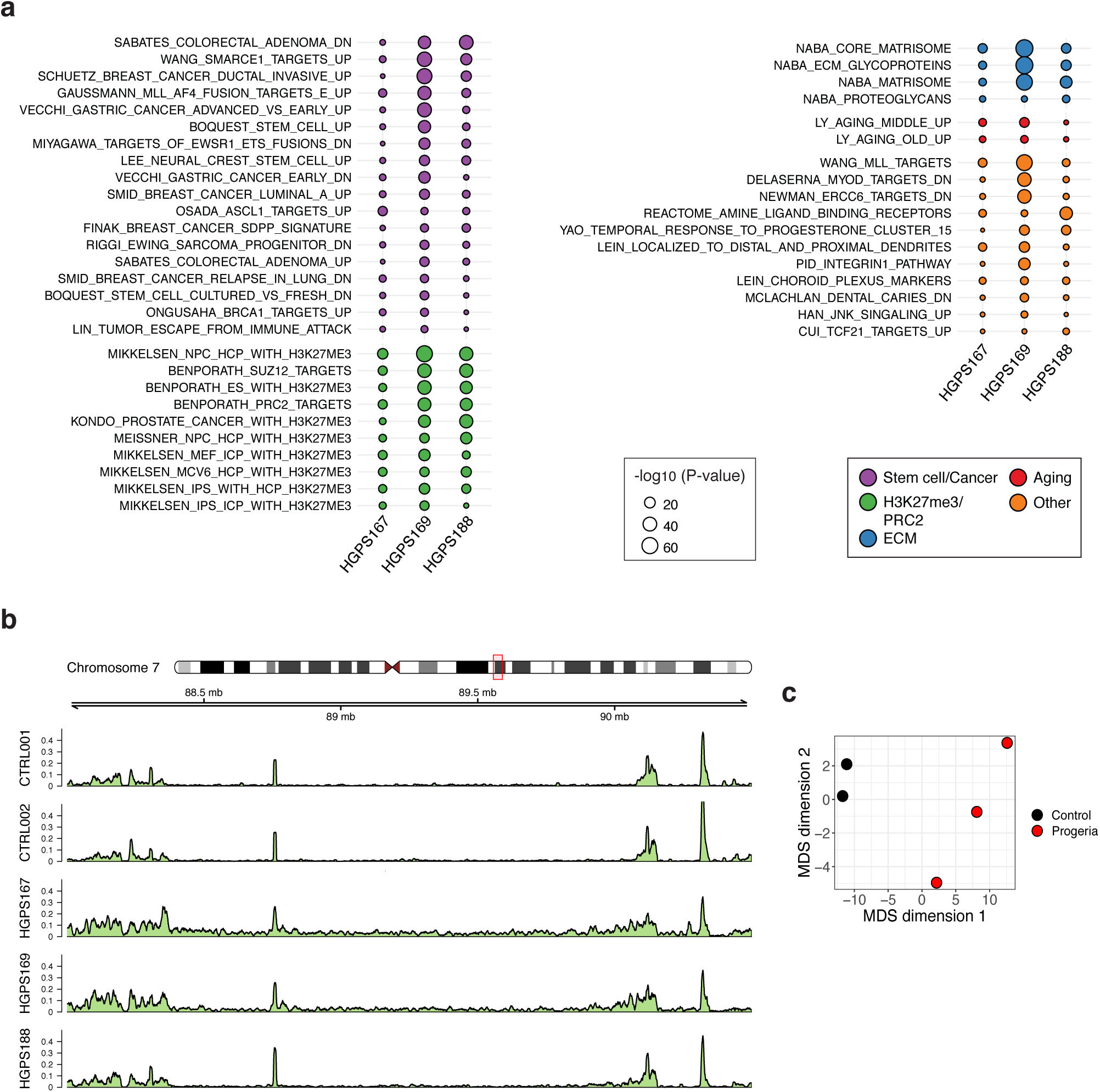
HGPS fibroblasts show an alteration of PRC2 distribution. **a**, MSigDB pathways consistently deregulated in all three progeria samples (FDR corrected aggregated p-value < 0.05), based on comparison of individual HGPS samples to the group of control samples (transcript level analysis - see Online methods). Pathways are ranked based on the sum of their -log_10_ transformed p-values across all 3 progeria samples. Dot colors highlight different categories of pathways as per graphical legend: related to stem cells or cancer, H3K27me3/PRC2 regulation, extracellular matrix, aging or other categories. Dot size is proportional to the -log_10_ transformed p-value. **b**, Genomic track for H3K27me3 ChIP-seq signal in 2 control and 3 HGPS samples for a representative region (2.5Mb region in chr7:88000000-90500000). Enrichment signals (computed using SPP) for each sample are shown. **c**, Multi-dimensional scaling of H3K27me3 ChIP-seq IP experiments using read counts in 10k genomic regions across all autosomes. Black dots show control, while red dots show progeria samples. The 2D distance of dots is representative of the pairwise distance of samples using the 10k genomic region read counts.

Finally, we performed ChIP-seq to dissect H3K27me3 distribution at the genome-wide level. By visual inspection we noticed that while the main peaks were present in both controls and HGPS samples, H3K27me3 signal was spread over flanking regions in HGPS (Fig. 5b). The progeria-associated differences in H3K27me3 profile were also confirmed by multi-dimensional scaling of ChIP-seq data (Fig. 5c), showing that control samples are close to each other and distant from the progeria samples, the latter also showing larger between sample distances. Analysis of TSS distribution of H3K27me3 mark revealed a slight decrease of signal in HGPS samples (Supplementary Fig. 6i). This difference was even more evident when considering the subset of upregulated genes in HGPS, where we observed a drop in H3K27me3 signal (Fig. 6a).

**Figure 6.**
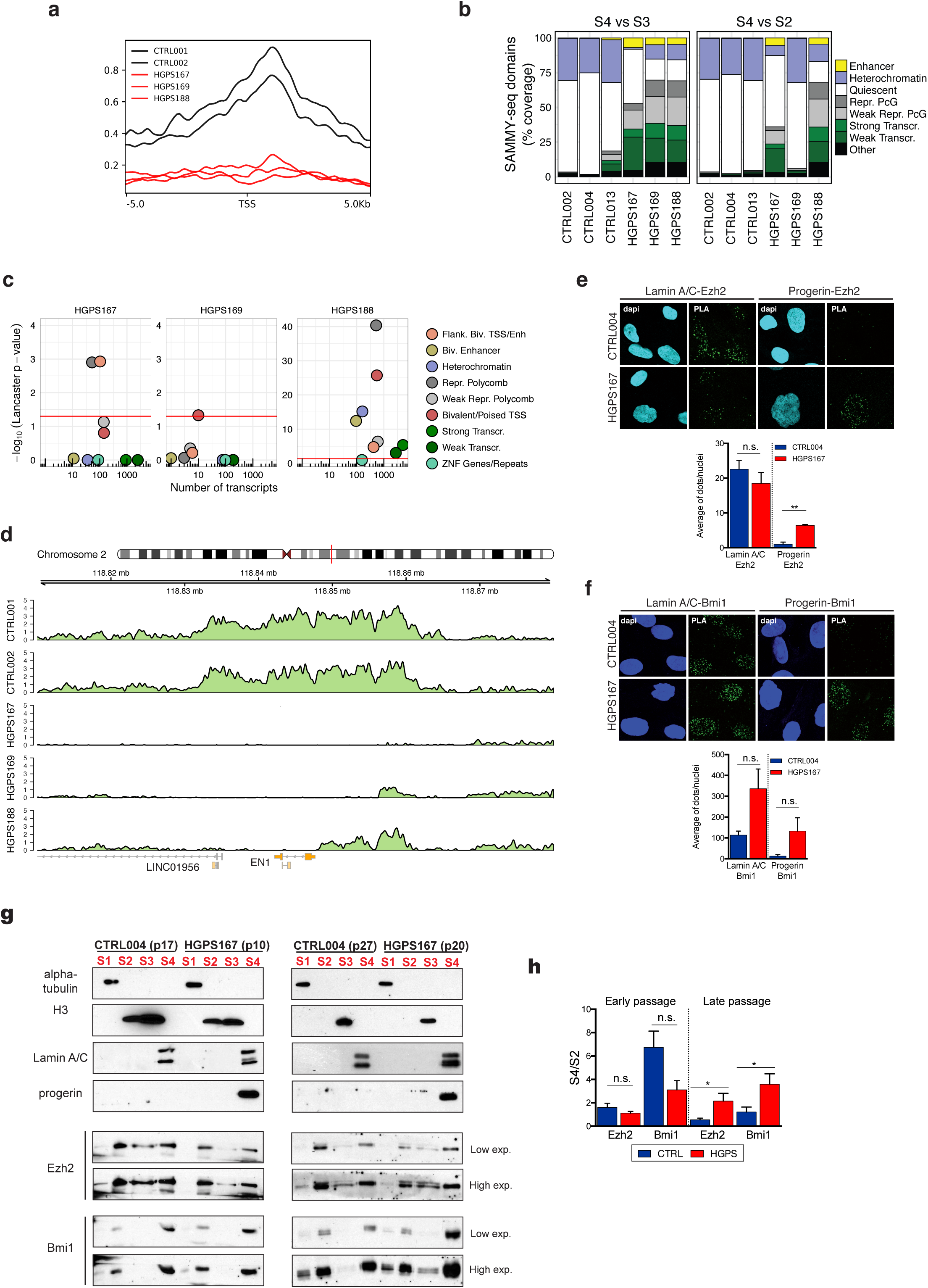
Chromatin structural changes in HGPS affect PRC2 regulated bivalent gens. **a**, H3K27me3 signal distribution around the TSS region of genes upregulated in HGPS samples compared to controls calculated by deepTools using the genome-wide signal from SPP (see Online methods for details). The x-axis represents relative genomic position around the TSS (+/− 5Kb), and the y-axis represents average signal intensity. **b**, Overlap of SAMMY-seq domains from each sample to Roadmap Epigenomics chromatin states for normal fibroblasts (E055). The overlaps with each chromatin state are reported as percentage over the total of SAMMY-seq domains (S4 vs S3 or S4 vs S2) for each sample. **c**, Transcripts differential expression (up-regulation) was assessed by comparing individual HGPS samples against the group of controls, then p-values were aggregated based on their chromatin state, considering only genes in SAMMY-seq domains (S4 vs S2) (see Online methods for details). The plot reports the number of transcripts (x-axis) and significance (Lancaster method aggregated p-values – y-axis) for each chromatin state (color legend) as defined by^43^. **d**, Representative track of H3K27me3 ChIP-seq signal in 2 control and 3 HGPS samples around the bivalent EN1 gene (chr2: 118810000-118880000), that was upregulated in HGPS samples. **e**, **f**, Representative images of PLA experiments in CTRL004 and HGPS167 at late passages (p18). Each fluorescent dot represents the co-localization of Lamin A/C or Progerin and Ezh2 (panel e) or Bmi1 (panel f). Nuclei were stained with dapi. All data were generated from an average of three independent experiments, whiskers represent SEM. **g**, Western blot on chromatin fractionation experiments of CTRL004 and HGPS167 at early (left) and late (right) passages. Equal amounts of each fraction were hybridized with indicated antibodies. Alpha-tubulin, histone H3 and Lamin A/C were used as loading controls respectively for S1, S2 and S3, S4. **h**, The graph shows quantifications of Ezh2 and Bmi1 in S4 fraction normalized on S2. Data points were generated from an average of at least two biological replicates, whiskers represent SEM. Comparisons were done using a two-tailed t-test in (e, f, h). Statistically significant differences are marked * p < 0.05; * * p < 0.01.

To further dissect the role of chromatin regulators in the disrupted expression patterns, we examined the overlap of SAMMY-seq domains with the chromatin states of normal skin fibroblasts^43^. SAMMY-seq domains of control samples are mostly overlapping “Quiescent” or “Heterochromatin” states (Fig. 6b), whereas SAMMY-seq domains of HGPS cells comprise regions transcribed or regulated by PcG proteins in normal skin fibroblasts. To analyze the correlation between gene expression, chromatin states and altered genome structure, we grouped SAMMY-seq domain genes on the basis of their chromatin state in normal skin fibroblasts and examined their expression difference between control and HGPS cells (Fig. 6c and Supplementary Fig. 6k). This analysis revealed that PcG regulated and bivalent genes are especially deregulated at the transcriptional level in progeria with respect to controls. These findings overall suggest that early chromatin remodelling has an impact on a subset of PcG targets, the bivalent genes, more susceptible to variations of PcG occupancy^49^ (Fig. 6d).

To verify the hypothesis of a PcG-progerin crosstalk we used the proximity ligation assay (PLA), which detects interactions between proteins in close proximity (< 30nm)^50^. We found that Lamin A interacts with PcG proteins in control or HGPS fibroblasts (Fig. 6 e, f), and that progerin, expressed in late-passage HGPS, interacts with both Ezh2 and Bmi1 (Fig. 6 e, f). To investigate the aberrant PcG-progerin interaction in HGPS progression we tested PcG protein compartmentalization, in early and late-passage HGPS (Fig. 6g, h). Comparing controls and early-passage HGPS fibroblasts we did not observe major differences in PcG nuclear distribution across chromatin fractions. On the other hand, PcG proteins were predominantly localized in the S4 fraction at late passage HGPS, supporting the previous observation that PcG regulated regions shift towards the insoluble S4 fraction in HGPS (Fig. 6b). Altogether, our results indicate that increased amount of progerin may alter PcG proteins nuclear compartmentalization and function.

## Discussion

Within the eukaryotic cell nucleus, chromatin acquires a specific structure which is fundamental for the correct spatiotemporal regulation of gene expression^51^. In particular, heterochromatin is characterized by highly compact structure and specific nuclear compartmentalization. Important heterochromatin organizing regions include LADs, along with pericentromeric and telomeric regions, which are crucial for preserving chromosome structure^3,4,10^. Alterations in heterochromatin are associated with developmental defects and cancer, while its proper conformation is a hallmark of healthy cells^52,53^. As such, reliable methods to characterize heterochromatin and its alterations are a crucial need for the biomedical scientific community.

Here we present a new technology called SAMMY-seq, for mapping lamina associated heterochromatic regions (Fig. 1). The protocol is based on the sequential extraction of multiple chromatin fractions, corresponding to increasingly compacted and less accessible chromatin regions, which are identified by the relative comparison of high-throughput sequencing reads from different fractions (Fig. 2). We show that SAMMY-seq can reproducibly and reliably identify lamina associated heterochromatic regions as the less accessible portions of chromatin in control wild-type fibroblasts. SAMMY-seq represents a significant improvement in the field of chromatin characterization as it provides novel information complementary and not overlapping with other high-throughput sequencing based methods commonly used to study chromatin structure and function. Of note, our protocol is overcoming several major limitations of other methods for mapping lamina associated heterochromatic regions. First of all, the procedure can be applied on primary cells, as it doesn’t require exogenous gene expression as in DamID-seq. Then, SAMMY-seq does not involve chemical modifications of chromatin, which might cause artifacts and high background after sequencing^54,55^. Moreover, it does not rely on antibodies for enriching specific chromatin fractions, thus avoiding issues related to antibody specificity, production, lot-variability and cross-reactivity. This is particularly important when studying chromatin changes in cells where protein levels of chromatin associated factors could be altered, thus allowing more flexibility in terms of experimental design compared to antibody-based techniques. Currently, we are successfully applying SAMMY-seq to many other experimental systems such as human epithelial and myogenic cells, human lymphocyte subpopulations or murine muscle satellite stem cells, in some cases scaling down to 10.000 cells (data not shown).

Emerging studies revealed that chromatin tethering at nuclear lamina and histone marks deposition are independent processes^56,57^ and that nuclear relocation is not sufficient to trigger a transcriptional switch^57–59^. Overall, our findings support these assumptions in HGPS model, further demonstrating that the pathological chromatin remodeling trigger a transcriptional dysfunction at PcG bivalent target, more sensitive to the epigenetic environment. In particular, by applying our SAMMY-seq technology on early passage HGPS fibroblasts we were able to detect chromatin structure alterations, in terms of accessibility changes (Fig. 3), despite the recent failure to do so by Hi-C^29^. Indeed Hi-C is designed to measure pairwise interactions between genomic loci, but they are not able to distinguish the intra-nuclear compartments where such interactions occur^60^. Notably, changes in chromatin accessibility seen in early-passage HGPS are not accompanied by H3K9me3 alterations or transcriptional deregulation in the same genomic regions (Fig. 4, Supplementary Fig. 5b, c), further indicating that chromatin remodeling detected by SAMMY-seq precedes H3K9me3 decrease observed at later passages (Fig. 4d). On the other hand, we found that chromatin accessibility changes in HGPS functionally affect PcG-regulated genes, by interfering with H3K27me3 genomic distribution (Fig. 5). In particular, the bivalent genes, involved in stemness maintenance and cell identity specification, are mostly affected in early-passage HGPS fibroblasts (Fig. 6), providing a mechanism for the previously postulated idea that the reduced potential for tissue homeostasis, commonly associated with progeria, is due to a misregulation of stem cell compartments^48^.

## Methods

### Cell cultures

Primary fibroblast cell lines were cultured in DMEM High glucose with glutamax supplemented with 15% FBS and 1% Pen/Strep. HGADFN164 (HGPS164), HGADFN167 (HGPS167), HGADFN169 (HGPS169), HGADFN188 (HGPS188), HGADFN271 (HGPS271) human dermal fibroblasts derived from HGPS patients were provided by the Progeria Research Foundation (PRF). AG08498 (CTRL001) and AG07095 (CTRL002) human dermal fibroblasts were obtained from the Coriell Institute. Preputial fibroblast strain #2294 (CTRL004) was a generous gift from the Laboratory of Molecular and Cell Biology, Istituto Dermopatico dell’Immacolata (IDI)-IRCCS, Rome, Italy”, while control dermal fibroblast CTRL013 was kindly provided by the Italian Laminopathies Network.

### Histochemistry, immunofluorescence assay and PLA analysis

Beta-galactosidase assay: cells were fixed at room temperature for 10min in paraformaldehyde at 4%, washed twice with 1×PBS, and incubated with fresh staining solution at 37 °C for 16 hrs in dark room. Then cells were washed twice with 1×PBS, overlaid with 70% glycerol/PBS, and photographed.

BrdU (5-Bromo-2¢-deoxy-uridine, Sigma B9285) labeling: cells were grown for 8 hrs in the presence of 50microM of BrdU, and then fixed in 4% PFA. After Triton X-100 treatment, cells were incubated for 2 min at RT in 0.07N NaOH, briefly rinsed twice in PBS and blocked in PBS/1% BSA. Reaction with BrdU antibody (1:10, Becton Dickinson 347580) was performed at room temperature for 1 hour in a PBS solution containing BSA 1%.

Immunofluorescence assay: coverslips were fixed with paraformaldehyde at 4% in PBS for 10 minutes. Then, cells were permeabilized with 0,5% triton X-100 in PBS and non-specific signals were blocked with 1% BSA in PBS for 30 minutes at room temperature. The following antibodies were used: Bmi1 (Millipore 05-637, mouse) diluted 1:100; Lamin A/C (Santa Cruz sc-6215, goat) diluted 1:200; Ezh2 (Cell signaling AC22 3147S, mouse) diluted 1:100; H3K9me3 (Abcam ab8898, rabbit) diluted 1:500; H3K27me3 (Millipore 07-449, rabbit) diluted 1:100. Incubation was performed at 12-16 hrs at 4°C or at room temperature for 2 hours for Lamin A/C. Primary antibodies were diluted in a PBS solution containing BSA 1%. Cells were stained with appropriate secondary antibodies, diluted 1:200 for 1 h at room temperature. Washes were done in PBT. As secondary antibodies we used Alexa Fluor 488 Donkey anti-mouse IgG (715-545-150), Alexa Fluor 594 Donkey anti-goat IgG (705-585-003) from Jackson ImmunoResearch Laboratories and Alexa Fluor 647 Chicken anti-goat IgG (Invitrogen, A21469). Finally, DNA was counterstained with DAPI, and glasses were mounted in Vectashield Antifade (Vector Laboratories) or ProLong Gold Antifade Reagent (Invitrogen). For PLA experiments, coverslips were fixed first with paraformaldehyde at 4% in PBS for 10 minutes. Then, cells were permeabilized with 0,5% triton X-100 in PBS and blocked with 1% BSA in PBS for 1 hour at room temperature. Incubation with progerin (Alexis human mAb, 13A4, ALX-804-662-R200) diluted 1:20 or Lamin A/C (Santa Cruz sc-6215, goat) diluted 1:200 was performed for 12-16 hrs at 4°C. After PBT washes, cells were incubated with Ezh2 (Cell signaling 4905S, rabbit 1:100) or Bmi1 (Abcam ab85688 rabbit 1:100) for 12-16 hrs at 4°C. Primary antibodies were diluted in a PBS solution containing BSA 1%. Finally, detection of protein interactions was performed using the Duolink system (Sigma) following manufacturer’s instructions.

### Image processing and analysis

Fluorescent images were taken with a Nikon ECLIPSE 90i microscope (1006 objective), equipped with a digital camera (Nikon Coolpix 990) and NIS Element software or with confocal Leica SP5 supported by LAS-AF software (version 2.6.0). Confocal image size was 1024⋅1024. PLA blobs quantification was performed using Cell Profiler 2.0 as previously described^18^. In order to automatically detect and quantify PcG bodies in fluorescence cell images we developed an algorithm that implements a method derived from a previous work ^65^, with variants and adaptations. The algorithm performs the segmentation of cell nuclei and the detection of PcG bodies of each image. It then measures the area and eccentricity of nuclei as well as the number and the area of the PcG bodies. It also provides for each PcG body a measure of its closeness to nuclear periphery (proximity). The algorithm has been implemented in MATLAB following this pseudocode:

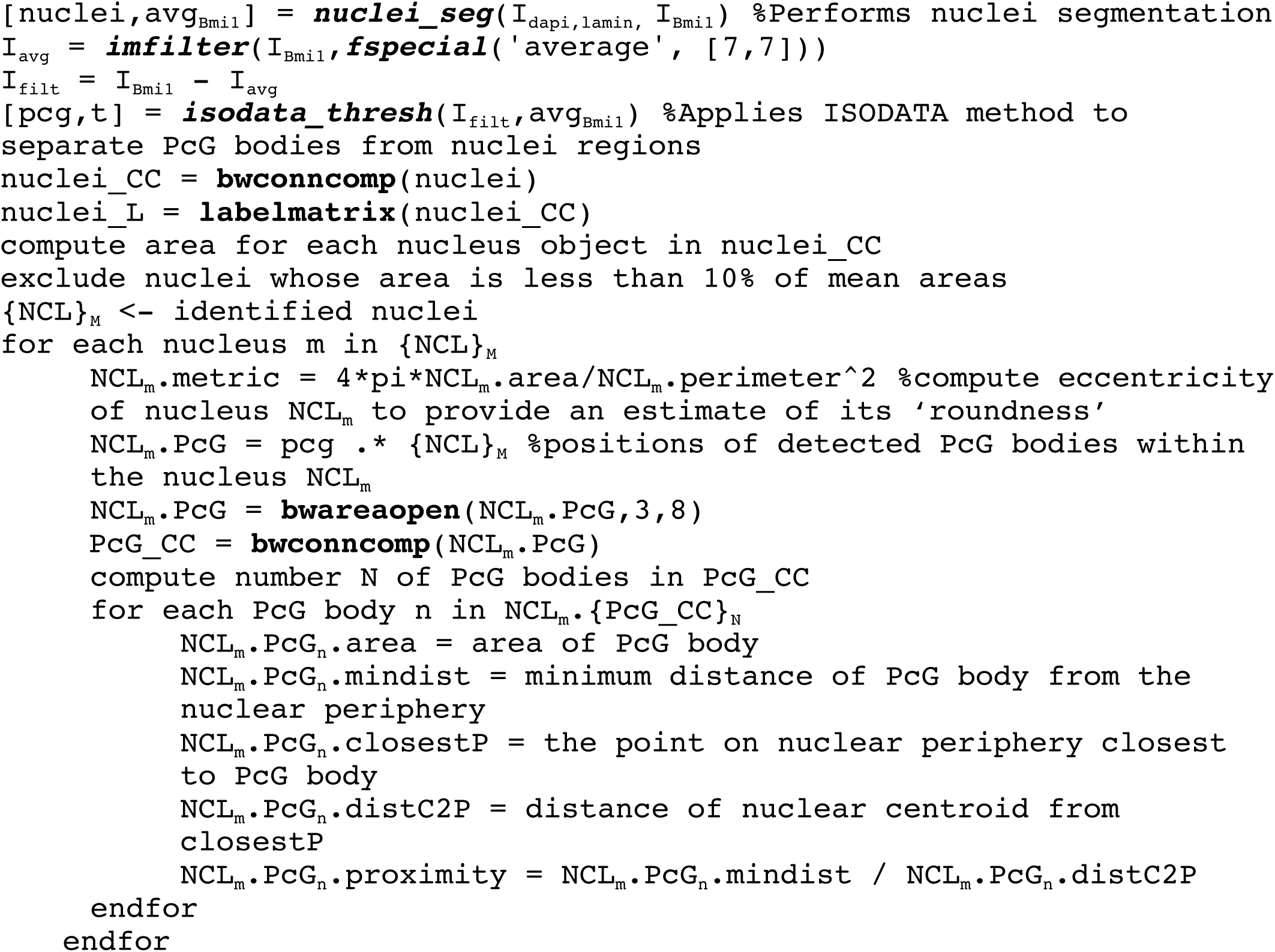

I_dapi,lamin_ is the image obtained by the sum of both images showing the fluorescence of nucleus and lamin.

The function *nuclei_seg* performs a partition of cell image I_dapi,lamin_ in nuclei regions and background implementing a region based segmentation algorithm^66^. `nuclei` is the binary image defining nuclei regions while avg_Bmi1_ is the mean intensity value of the nuclei regions in the image I_Bmi1_ that shows the fluorescence of PcG bodies.

In order to better enhance PcG areas we subtract from the original image I_Bmi1_ its smoothed version obtained by applying an averaging filter of size 7, producing the image I_filt_.

The function *isodata_thresh* implements the ISODATA classification algorithm^67^ and uses relevant values computed by *nuclei_seg* function in order to extract PcG bodies from the nuclei regions. It sets the initial threshold value of ISODATA method as avg_Bmi1_. pcg is the binary image defining PcG bodies.

Reconstructions of nuclei are obtained through the connected components algorithm (*bwconncomp* MATLAB function, using a connectivity of 6). Nuclei are then labeled by applying the *labelmatrix* MATLAB function so they can be easily separated from each other.

The algorithm computes the area of each nucleus, discarding objects whose area is less than 10% of mean areas which are just noise.

The algorithm uses the *bwareaopen* function in order to discard too small detected PcG objects which are probably just noise.

Reconstructions of PcG bodies are obtained through the connected components algorithm (*bwconncomp* MATLAB function, using a connectivity of 6).

The algorithm computes the number of PcG bodies, the area of any PcG body, and the minimum euclidean distance of any PcG body from the nuclear periphery.

### Protein extraction and Western Blot analyses

Total proteins were prepared by resuspending 1×10^6^ cells in extraction buffer (50 mM TrisHCl pH 7.6; 0.15 M NaCl; 5 mM EDTA; 16 Protease Inhibitors; 1% Triton X-100). One 40sec pulse of sonication (UP100H manual sonicator, Hielscher) at 40% amplitude was performed to allow dissociation of protein from chromatin and solubilization. Extracts were analyzed by SDS-PAGE using an 8% gel (37.5:1 Acryl/Bis Acrylamide). The following primary antibodies were used: Beta-Actin (Santa-Cruz sc1616, rabbit 1:4000), H3 total (Abcam ab1791, rabbit 1:6000), Lamin A/C (Santa Cruz sc-6215, goat 1:4000), Lamin B (Santa Cruz sc6216, goat 1:2000), progerin (13A4 mouse, Abcam 66587, mouse 1:1000), Ezh2 (AC22 Cell Signaling 3147S, mouse 1:1000), Bmi1 (D42B3 Cell signaling, rabbit 1:1000), H3K9me3 (Abcam ab8898, rabbit 1:1000), H3K27me3 (Millipore 07-449 rabbit 1:1000).HRP-conjugated secondary antibodies were revealed with the ECL chemiluminescence kit (Thermo Fisher Scientific).

### Chromatin fractionation

Chromatin fractionation was carried out as described in^38^ with minor adaptions. Briefly, 4 million fibroblasts were washed in PBS 1X, and extracted in cytoskeleton buffer (CSK: 10 mM PIPES pH 6,8; 100 mM NaCl; 1 mM EGTA; 300 mM Sucrose; 3 mM MgCl_2_; 1X protease Inhibitors by Roche Diagnostics; 1 mM PMSF) supplemented with 1 mM DTT and 0,5% Triton X-100. After 5 min at 4°C the cytoskeletal structure was separated from soluble proteins by centrifugation at 3000 rpm for 3 min, and the supernatant was labeled as S1 fraction. The pellets were washed with an additional volume of cytoskeleton buffer. Chromatin was solubilized by DNA digestion with 25U of RNase–free DNase (Turbo DNAse; Invitrogen AM2238) in CSK buffer for 60 min at 37°C. To stop digestion, ammonium sulphate was added in CSK buffer to a final concentration of 250 mM and, after 5 min at 4°C samples were pelleted at 5000 rpm for 3 min at 4°C and the supernatant was labeled as S2 fraction. After a wash in CSK buffer, the pellet was further extracted with 2M NaCl in CSK buffer for 5 min at 4°C, centrifuged at 5000 rpm 3 min at 4°C and the supernatant was labeled as S3 fraction. This treatment removed the majority of histones from chromatin. After 2 washing in NaCl 2M CSK, the pellets were solubilized in 8M urea buffer to remove any remaining protein component by applying highly denaturing conditions. This remaining fraction was labeled as S4. Supernatants from each extraction step were quantified and analyzed by SDS-PAGE and immunoblotting. Anti-tubulin alpha (Sigma T5168, mouse 1:10000), H3 (Abcam ab1791, rabbit 1:4000), Beta-Actin (Santa-Cruz sc1616, rabbit 1:4000), Lamin A/C (Santa Cruz sc-6215, goat 1:4000), Lamin B (Santa Cruz sc-6216, goat 1:2000), progerin (13A4 mouse, Abcam 66587, mouse 1:1000), Ezh2 (AC22 Cell Signaling 3147S, mouse 1:1000), Bmi1 (D42B3 Cell signaling, rabbit 1:1000) were used as primary antibodies. HRP-conjugated secondary antibodies were revealed with the ECL chemiluminescence kit (Thermo Fisher Scientific).

### DNA sonication and sequencing for chromatin fractionation

For DNA fractionation, we took an aliquot corresponding to 30% of volume of S2, S3, S4 fractions and added TE buffer to reach a final volume of 300µL. After incubation with RNAse A (Roche) (90 minutes at 37°) and Proteinase K (Sigma) (150 minutes at 55°), DNA was extracted from beads by standard phenol/chloroform extraction, precipitated and resuspended in 25µl milliQ H_2_O. After Nanodrop (260/280 = 1,7-1,9; 260/230 ≥ 2) and Qubit HS DNA quantification, we added H_2_O to a final volume of 105µL. Then samples were transferred to 96 well plates and sonicated 4 times with Bioruptor sonicator (10 minutes 30 seconds ON - 30 seconds OFF, High Power). The DNA profiles were finally checked by capillary electrophoresis (Agilent 2100 Bioanalyzer). Finally, DNA libraries were prepared by using NuGEN Ovation Ultralow Library Prep System kit and then sequenced using an Illumina HiSeq 2500 instrument according to manufacturer’s instructions (Illumina).

### Chromatin immunoprecipitation and sequencing

Cells were cross-linked with 1% HCHO for 12 minutes at room temperature, lysed and chromatin sheared. IP were performed overnight on a wheel at 4° with 2.4μg of H3K9me3 antibody (ab8898, Abcam) or 2μg of H3K27me3 (07-449, Millipore). The following day, antibody-chromatin immunocomplexes were loaded onto Dynabeads Protein G (Invitrogen 10004D). The bound complexes were washed once in Low Salt Solution (0,1% SDS, 2mM EDTA, 1% Triton X-100, 20mM Tris pH 8, 150 mM NaCl), once in High Salt Solution (0,1% SDS, 2mM EDTA, 1% Triton X-100, 20mM Tris pH 8, 500 mM NaCl), once again in Low Salt Solution and once in Tris/EDTA 50 mM NaCl. Crosslinking was reversed at 65°C overnight in Elution Buffer (50mM Tris pH 8, 20mM EDTA, 1%SDS), DNA was purified by standard phenol/chloroform extraction, precipitated and resuspended in 30µl of 10mM Tris pH 8. ChIP efficiency was tested by qPCR reactions, performed in triplicate using QuantiTect SYBR Green master mix (Qiagen) on a StepOnePlus™ Real-Time PCR System (Applied Biosystems). Relative enrichment was calculated as IP / Input ratio. Libraries for sequencing were created using the automation instrument Biomek FX (Beckman Coulter), then qualitatively and quantitatively checked using Agilent High Sensitivity DNA Kit (Agilent Technologies, 5067-4627) on a Bioanalyzer 2100 (Agilent Technologies). Libraries with distinct adapter indexes were multiplexed and, after cluster generation on FlowCell, were sequenced for 50bp in the single read mode on a HiSeq 2000 sequencer at the IEO Genomic Unit in Milan.

### RNA sequencing

For high-throughput sequencing, cDNA libraries were prepared from total RNA, extracted with Trizol, by using Illumina TruSeq Stranded Total RNA Kit with Ribo-Zero GOLD. cDNA fragments of ~300 bp were purified from each library and were sequenced for 125bp, using an Illumina HiSeq 2500 instrument according to manufacturer’s instructions (Illumina).

### SAMMY-seq sequencing read analysis

Sequencing reads were aligned to the hg38-noalt reference human genome available in the bcbio-nextgen pipeline, using bwa aln^61^ (version 0.7.12) with options -n 2 -k 2 and saved the results in sam format with bwa samse. The sam files were converted to bam and name sorted with samtools^62^ (version 1.3.1). We marked PCR duplicates using the biobambam2 toolset^63^ (version 2.0.54). We discarded reads mapping to non-autosomal chromosomes, PCR duplicates, qcfail, multimapping and unmapped reads with samtools. We converted the bam files to bedgraph using bedtools^64^ (version 2.25.0) and bedgraph to bigWig using UCSC’s bedgraphToBigWig tool (version 4) for reads distribution visualization. Read coverage was normalized by the total sequencing library size, before converting bedgraph files to bigWig. Sample quality was assessed using fastqc (version 0.11.5) (http://www.bioinformatics.babraham.ac.uk/projects/fastqc). For the down sampling analysis, we sampled the raw fastq files to 50% and 25% using the seqtk sample command (https://github.com/lh3/seqtk) and ran the same pipeline, as above.

### ChIP-seq sequencing read analysis

Sequencing reads were aligned, filtered, converted and quality checked using the same tools as for the SAMMY-seq reads. Before sequence alignment we did an additional trimming step, using Trimmomatic^65^ (version 0.32) in single end mode for the H3K9me3 data. We used the Enriched Domain Detector (EDD)^40^ tool to call H3K9me3 peaks, with the following options: --fdr 0.1 –gap-penalty 10 –bin-size 100 –write-log-ratios –write-bin-scores and also excluding blacklisted genomic regions containing telomeric, centromeric, and certain heterochromatic regions^66^. We also changed the required_fraction_of_informative_bins parameter to 0.98. We calculated genome wide ChIP-seq signal for H3K9me3 and H3K27me3 data, using the SPP package (version 1.15.4)^67^. We imported bam files into the R (version 3.3.1) statistical environment, and selected informative reads with the get.binding.characteristics and select.iformative.tags functions, removed anomalous positions with extremely high number of reads using the remove.tag.anomalies function, and calculated the differential signal, smoothed by a Gaussian kernel, using the get.smoothed.tag.density function with the default bandwidth and tag.shift parameters. In the case of H3K27me3 data, we also set the scale.by.dataset.size = TRUE parameter for the get.smoothed.tag.density function.

MDS analysis was done using the plotMDS function of edgeR (version 3.24.3)^68^ after importing the 10k genomic region read counts for all samples into the R (version 3.5.1) statistical environment. We dropped all genomic regions where the log count-per-million (cpm) value did not reach at least 1 in at least 2 samples. We calculated the normalization factor of samples using the calcNormFactors and estimated dispersion using the estimateDisp function of edgeR. We used the top = 1000 and gene.selection = “common” parameters for the plotMDS function. The gene.selection parameter here does not refer to genes, but the imported genomic regions. We used deepTools (version 3.2.1) ^69^ to visualize the ChIP-seq signal around TSS regions, with the following parameters: --referencePoint TSS -b 5000 -a 5000 –missingDataAsZero – averageTypeBins median –binSize 100.

### Literature data processing

We collected publicly available datasets from the following sources: ATAC-seq from^45^, Lamin A/C ChIP-seq from^29^, Lamin B ChIP-seq from^44^. Sequencing reads were aligned, filtered and converted using the same tools as for the SAMMY-seq reads. We calculated the genome wide ChIP-seq signal for Lamin A/C and Lamin B data, using the SPP package (version 1.15.4)^67^. We imported bam files into the R (version 3.3.1) statistical environment, and selected informative reads with the get.binding.characteristics and select.informative.tags functions, removed anomalous positions with extremely high number of reads using the remove.local.tag.anomalies function, and calculated the differential signal, smoothed by a Gaussian kernel, using the get.smoothed.tag.density function with the default bandwidth and tag.shift parameter. We used the filtered read count as the genome-wide signal for the ATAC-seq sample.

Additionally, we downloaded from Roadmap Epigenomics^43^ the genome-wide signal coverage tracks for the DNAse-seq, as well as histone marks ChIP-seq datasets (H3K9me3, H3K27me3, H3K4me1, H3K36me3, H3K27ac, H3K4me3) for the foreskin fibroblast sample E055. We converted bigwig files to bedgraph with the bigWigToBedGraph tool, lifted over genomic coordinates from hg19 to hg38 with the liftOver tool and converted back bedgraph to bigwig with the bedGraphToBigWig tool using the UCSC toolkit^70^ (version 4).

We calculated genome-wide correlations between SAMMY-seq samples and all of the public datasets listed and processed above using StereoGene^46^ (version 1.73). StereoGene uses kernel correlation to calculate genome-wide correlation for spatially related but not completely overlapping features irrespective of their discrete or continuous nature.

We called lamina associated domains with the EDD tool. For the Lamin A/C dataset we used the following options: –gap-penalty 25 –bin-size 200 –write-log-ratios –write-bin-scores, and for the Lamin B dataset we used: –gap-penalty 5 –bin-size 100 –write-log-ratios –write-bin-scores. We also changed the required_fraction_of_informative_bins parameter to 0.98.

### SAMMY-seq domain calling, signal calculation, overlap analysis and visualization

We performed relative comparisons of SAMMY-seq fractions within each sample using EDD, optimized to call very broad enrichment domains. EDD was originally designed for lamin ChIP-seq data, comparing IP and input samples. As SAMMY-seq data also shows broad enrichment regions, we used EDD to select significantly enriched SAMMY-seq domains by comparing less accessible fractions to more accessible ones (S3 vs S2, S4 vs S3 and S4 vs S2 comparisons) in each sample, with the following options: --gap-penalty 25 --bin-size 50 --write-log-ratios --write-bin-scores and also excluding blacklisted genomic regions containing telomeric, centromeric, and certain heterochromatic regions^66^. We also changed the required_fraction_of_informative_bins parameter to 0.98. We used the same set of parameters for the down sampled data, except changing –bin-size to 100 for 50% down sampling or 200 for 25% down sampling.

Additionally, we calculated the genome wide differential signal for all comparisons, using the SPP package (version 1.15.4)^67^. We imported bam files into the R (version 3.3.1) statistical environment, and selected informative reads with the get.binding.characteristics and select.informative.tags functions, removed anomalous positions with extremely high number of reads using the remove.local.tag.anomalies function, and calculated the differential signal, smoothed by a Gaussian kernel, using the get.smoothed.tag.density function with the default bandwidth parameter and tag.shift = 0.

We used bedtools^64^ (version 2.25.0) and the bedtools jaccard or bedtools fisher command to calculate Jaccard Index or Fisher-test p-values for the various overlap analyses. We randomized SAMMY-seq domains and LADs for the SAMMY-seq – LAD overlap analysis using the bedtools shuffle with -noOverlapping option and also excluding blacklisted genomic regions containing telomeric, centromeric, and certain heterochromatic regions^66^ with the -exclude option.

We used the gViz^71^ and karyoploter^72^ Bioconductor packages to visualize SAMMY-seq read coverages, differential signals and domains. We used ggplot2^73^ for additional plotting, and the GAM method^74^ from the mgcv package (version 1.8-12) for smoothing data while plotting SAMMY-seq border region signals.

### RNA-seq sequencing read analysis

Transcript and gene level quantification was done with Kallisto^75^ (version 0.43.0) to estimate transcript level read counts and TPM (Transcripts Per Million) values. The GENCODE v27 annotation was used to build the transcriptome index. Kallisto was run with the --bias option to perform sequence based bias correction on each sample. We used sleuth^76^ (version 0.29.0) to calculate gene or transcript level differential expression comparing all three controls against all three HGPS samples, where the linear model used in sleuth included the sample type (control or HGPS), sex and library id. Additionally, we analyzed transcript level differential expression, comparing the three controls with a single progeria sample, where the linear model used in sleuth included the sample type (control or HGPS) and sex. For both cases, we used sleuth’s Wald test to calculate p and q-values. After differential expression, we grouped transcripts according to MSigDB pathways and aggregated transcript p-values using the Lancaster aggregation method from the aggregation R package (version 1.0.1) motivated by a recently described analysis pipeline in^77^. Lancaster p-values were corrected for multiple testing using the Benjamini-Hochberg method. Alternatively, we aggregated transcript p-values based on their Roadmap chromatin states and differential expression direction. We used two different criteria: for the Tx, TxWk, ReprPC, ReprPCWk, Het, and ZNF/Rpts chromatin states, we filtered for transcripts overlapping at least 50% with the state. For the EnhBiv, TssBiv and BivFlnk regions we required that the 200nt region around the TSS region overlaps at least 50% with the state. We aggregated p-values of up-regulated (sleuth b-value > 0) or down-regulated (sleuth b-value < 0) transcripts separately, according to their chromatin state overlap and additionally filtering for being in a specific (S4 vs S3 or S4 vs S2) enrichment region.

## Data availability

The datasets generated during the current study are available in the GEO repository with accession number PRJNA483177.

## Acknowledgements

We thank Giovanna Lattanzi, Sammy Basso, the Italian network of Laminopathies and members of the laboratory for stimulating discussions and constructive criticisms. We thank Beatrice Bodega, Marina Lusic, Maria Vivo, Daniela Palacios, Chiara Mozzetta, Judith Hariprakash, Koustav Pal, Paolo Maiuri, Mattia Forcato and Fabrizio d’Adda di Fagagna for critical feedback on the manuscript. We are grateful to Mariangela Panetta and Valentina Saccone for providing support with cell cultures. We thank Elisa Salviato for advice on statistical analyses. We gratefully acknowledge the Progeria Research Foundation for providing primary human fibroblasts of HGPS patients. This work was supported by grants from the flagship CNR projects, (Epigen and Interomics) to C.L. and G.O, the Italian Ministry of Health n. GR-2013-02355413 to C.L., My First AIRC Grant (MFAG) n. 18535 to C.L., AFM-Telethon n. 21030 to C.L. and F.F.; by AIRC Start-up grant 2015 n.16841 to F.F.; and Cariplo 2017-0649 to C.L. and F.F. E.S. was supported by the Structured International Post-doc Program of SEMM (SIPOD) and the AFM-TELETHON fellowship n. 21835.

## Author Contributions

C. L. designed and F. M., F. L., A. B. and I.O. performed the experiments. E. S., C. P. and F. F. analyzed all sequencing data. S. V. contributed to analyze ChIP-seq data. L.A, F.G and G.O performed image processing and analysis. E. S., F. M., C. L. and F. F. interpreted the results and wrote the manuscript. All authors reviewed and approved the manuscript for submission.

