## Supplementary Material for "Early Polycomb-target deregulations in Hutchinson-Gilford Progeria Syndrome revealed by heterochromatin analysis"

| Sample ID | Fraction | Read number |
|---|---|---|
| CTRL002 | S2 | 78102212 |
| CTRL002 | S3 | 61341611 |
| CTRL002 | S4 | 55731571 |
| CTRL004 | S2 | 78683296 |
| CTRL004 | S3 | 60438514 |
| CTRL004 | S4 | 54864540 |
| CTRL013 | S2 | 59635985 |
| CTRL013 | S3 | 65360683 |
| CTRL013 | S4 | 85335120 |
| HGPS167 | S2 | 58983114 |
| HGPS167 | S3 | 73501327 |
| HGPS167 | S4 | 61985833 |
| HGPS169 | S2 | 79209359 |
| HGPS169 | S3 | 77731616 |
| HGPS169 | S4 | 92435639 |
| HGPS188 | S2 | 68067662 |
| HGPS188 | S3 | 104711260 |
| HGPS188 | S4 | 54037646 |

**Supplementary Table 1.** Total number of sequencing reads in all samples and fractions for the SAMMY-seq sequencing runs.

| Sample ID | Comparison | Total length | Domain count | Domain average length | Autosome % |
|---|---|---|---|---|---|
| CTRL002 | S3 vs S2 | 605,736895 | 273 | 2,218816465 | 21,06909824 |
| CTRL002 | S4 vs S2 | 616,186907 | 285 | 2,162059323 | 21,43257672 |
| CTRL002 | S4 vs S3 | 582,086898 | 276 | 2,1090105 | 20,24649008 |
| CTRL004 | S3 vs S2 | 374,950204 | 204 | 1,837991196 | 13,04173932 |
| CTRL004 | S4 vs S2 | 499,250205 | 205 | 2,435366854 | 17,36521533 |
| CTRL004 | S4 vs S3 | 477,000188 | 188 | 2,537235043 | 16,5913021 |
| CTRL013 | S3 vs S2 | 644,500193 | 193 | 3,339379238 | 22,41738615 |
| CTRL013 | S4 vs S2 | 510,136889 | 267 | 1,910625052 | 17,7438824 |
| CTRL013 | S4 vs S3 | 435,23686 | 238 | 1,828726303 | 15,13866538 |
| HGPS167 | S3 vs S2 | 178,500104 | 104 | 1,716347154 | 6,208695983 |
| HGPS167 | S4 vs S2 | 549,050231 | 231 | 2,376840827 | 19,09738923 |
| HGPS167 | S4 vs S3 | 487,250195 | 195 | 2,498718949 | 16,9478239 |
| HGPS169 | S3 vs S2 | 782,150186 | 186 | 4,205108527 | 27,20520946 |
| HGPS169 | S4 vs S2 | 465,712191 | 260 | 1,791200735 | 16,19867633 |
| HGPS169 | S4 vs S3 | 629,494655 | 176 | 3,576674176 | 21,89545467 |
| HGPS188 | S3 vs S2 | 540,336884 | 262 | 2,062354519 | 18,79431645 |
| HGPS188 | S4 vs S2 | 650,582792 | 197 | 3,302450721 | 22,6289547 |
| HGPS188 | S4 vs S3 | 693,286821 | 199 | 3,483853372 | 24,11431144 |

**Supplementary Table 2.** Characteristics of SAMMY-seq domains, detected by EDD. The table contains the following columns. *Sample ID*: patient ID, *Comparison*: fractions used in the EDD run as the “IP” and “input” sample, *Total length*: total length of SAMMY-seq domains in megabase, *Domain count*: number of SAMMY-seq domains, *Domain average length*: average length of SAMMY-seq domains in megabase, *Autosome %*: % of autosomes covered by SAMMY-seq domains.

| Sample ID | Comparison | Lamin A overlap JI | Lamin A overlap % | Lamin B overlap JI | Lamin B overlap % |
|---|---|---|---|---|---|
| CTRL002 | S3 vs S2 | 0,609681 | 79,95224036 | 0,437656 | 60,95883699 |
| CTRL002 | S4 vs S2 | 0,598586 | 78,33665691 | 0,439887 | 60,6553253 |
| CTRL002 | S4 vs S3 | 0,584696 | 79,55001584 | 0,43488 | 61,92377326 |
| CTRL004 | S3 vs S2 | 0,406604 | 80,78410455 | 0,292558 | 59,28790505 |
| CTRL004 | S4 vs S2 | 0,58281 | 86,44967451 | 0,363631 | 59,09864797 |
| CTRL004 | S4 vs S3 | 0,603878 | 90,76519916 | 0,356267 | 59,70649895 |
| CTRL013 | S3 vs S2 | 0,630849 | 79,06904577 | 0,38012 | 53,49107836 |
| CTRL013 | S4 vs S2 | 0,502858 | 77,59685993 | 0,367625 | 58,87638469 |
| CTRL013 | S4 vs S3 | 0,31422 | 60,87493253 | 0,277669 | 52,05214556 |

**Supplementary Table 3.** Statistics on on control sample SAMMY-seq domains overlapping Lamina Associated Domains (LADs) based on literature data. The table contains the following columns. *Sample ID*: patient ID, *Comparison*: fractions used in the EDD run as the “IP” and “input” sample, *Lamin A overlap JI*: Jaccard index of the SAMMY-seq domain – Lamin A/C LAD comparison as calculated by bedtools, *Lamin A overlap %*: % of SAMMY-seq domains overlapping with Lamin A/C LADs, *Lamin B overlap JI*: Jaccard index of the SAMMY-seq domain – Lamin B LAD comparison as calculated by bedtools, *Lamin B overlap %*: % of SAMMY-seq domains overlapping with Lamin B LADs.

| Sample ID 1 | Sample ID 2 | Comparison | Overlap JI | Overlap number |
|---|---|---|---|---|
| CTRL002 | CTRL004 | S3 vs S2 | 0,536316 | 177 |
| CTRL002 | CTRL013 | S3 vs S2 | 0,585113 | 188 |
| CTRL004 | CTRL013 | S3 vs S2 | 0,39536 | 137 |
| HGPS167 | HGPS169 | S3 vs S2 | 0,190102 | 84 |
| HGPS167 | HGPS188 | S3 vs S2 | 0,282751 | 93 |
| HGPS169 | HGPS188 | S3 vs S2 | 0,482547 | 186 |
| CTRL002 | CTRL004 | S4 vs S3 | 0,686345 | 187 |
| CTRL002 | CTRL013 | S4 vs S3 | 0,46086 | 178 |
| CTRL004 | CTRL013 | S4 vs S3 | 0,370164 | 127 |
| HGPS167 | HGPS169 | S4 vs S3 | 0,172627 | 107 |
| HGPS167 | HGPS188 | S4 vs S3 | 0,121931 | 100 |
| HGPS169 | HGPS188 | S4 vs S3 | 0,717687 | 172 |
| CTRL002 | CTRL004 | S4 vs S2 | 0,707946 | 200 |
| CTRL002 | CTRL013 | S4 vs S2 | 0,695666 | 242 |
| CTRL004 | CTRL013 | S4 vs S2 | 0,614278 | 190 |
| HGPS167 | HGPS169 | S4 vs S2 | 0,168257 | 76 |
| HGPS167 | HGPS188 | S4 vs S2 | 0,0748601 | 85 |
| HGPS169 | HGPS188 | S4 vs S2 | 0,0823542 | 71 |

**Supplementary Table 4.** Pairwise comparison of SAMMY-seq domains across control or progeria samples. The table contains the following columns. *Sample ID 1*: patient ID for comparison, *Sample ID 2*: patient ID for comparison, *Comparison*: fractions used in the EDD run as the “IP” and “input” sample, *Overlap JI*: Jaccard index of the SAMMY-seq domain overlaps as calculated by bedtools, *Overlap number*: number of overlapping SAMMY-seq domains.

### Supplementary Figure 1

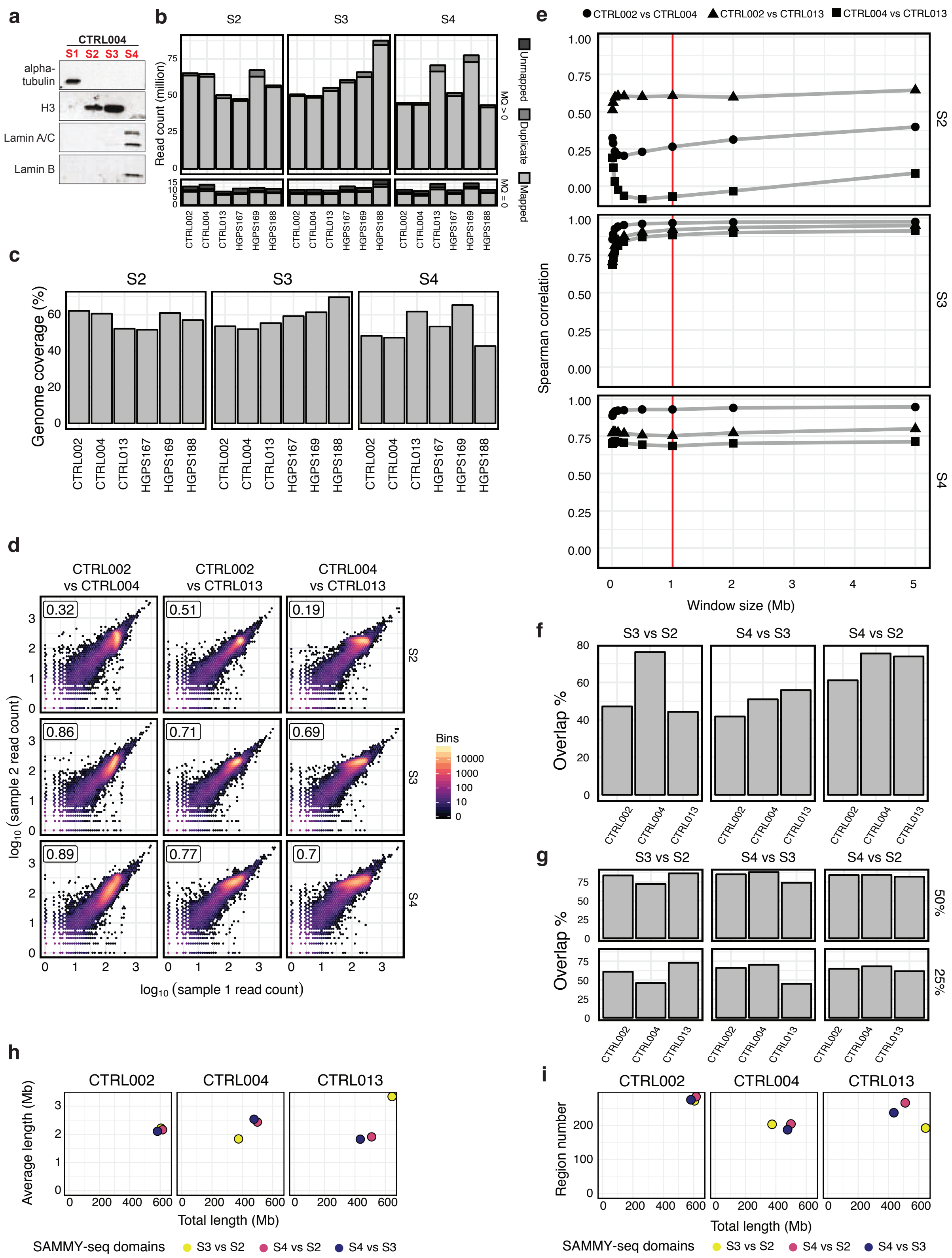

**Supplementary Figure 1 - Characteristics and reliability of SAMMY-seq libraries. a**, Western blots for chromatin fractionation experiments of CTRL004. Sequential extractions were performed to isolate soluble proteins (S1-fraction), DNase-sensitive chromatin (S2-fraction), DNase-resistant chromatin (S3-fraction), and the most compact and inaccessible chromatin (S4-fraction). Equal amounts of each fraction were hybridized with indicated antibodies. **b**, Total read counts for each SAMMY-seq sample. The stacked bar plot shows mapped, unmapped, and PCR duplicate reads with bwa mapping quality equal to zero (MQ = 0, non-unique mapping) or larger than zero (MQ > 0, uniquely mapped reads). The average number of sequencing reads in the different fractions were 72 million (S2), 62 million (S3) and 65 (S4) million. **c**, Percent of the human genome (hg38-noalt build) covered by at least one read in each SAMMY-seq sample. On average 58% (S2), 54% (S3) and 52% (S4) of the genome was covered by at least one read. **d**, Scatter plots comparing read counts over 10Kb bins in the S2, S3 and S4 fractions (rows) across three pairwise comparisons of the control samples (columns). The color gradient represents the number of genomic bins with the same (x,y) values. Spearman correlation values are reported in the top left corner of each subpanel. **e**, Spearman correlation (y-axis) of read counts between control samples pairs (shape of dots) over different bin sizes (x-axis) in the S2, S3 and S4 fractions. The red line shows the 1Mb bin size. **f**, Percentage of SAMMY-seq domains (S3 vs S2, S4 vs S3 and S4 vs S2) conserved across all 3 control samples (computed as percent over per sample total SAMMY-seq domains size). **g**, Percentage of SAMMY-seq domains (S3 vs S2, S4 vs S3 and S4 vs S2) detected after 50% or 25% down sampling of sequencing reads (computed as percent over per sample total SAMMY-seq domains size detected with 100% of sequencing reads). **h**, Average (y-axis) and total (x-axis) size of SAMMY-seq domains (S3 vs S2, S4 vs S3 and S4 vs S2) for each control sample. Average size across samples is 2.47Mb for S3 vs S2, 2.16Mb for S4 vs S3 and 2.17Mb for S4 vs S2. Average genome coverage across samples is 18.84% S3 vs S2, 17.33% for S4 vs S3 and 18.85% for S4 vs S2. **i**, Number (y-axis) and total size (x-axis) of SAMMY-seq domains (S3 vs S2, S4 vs S3 and S4 vs S2) for each control sample.

### Supplementary Figure 2

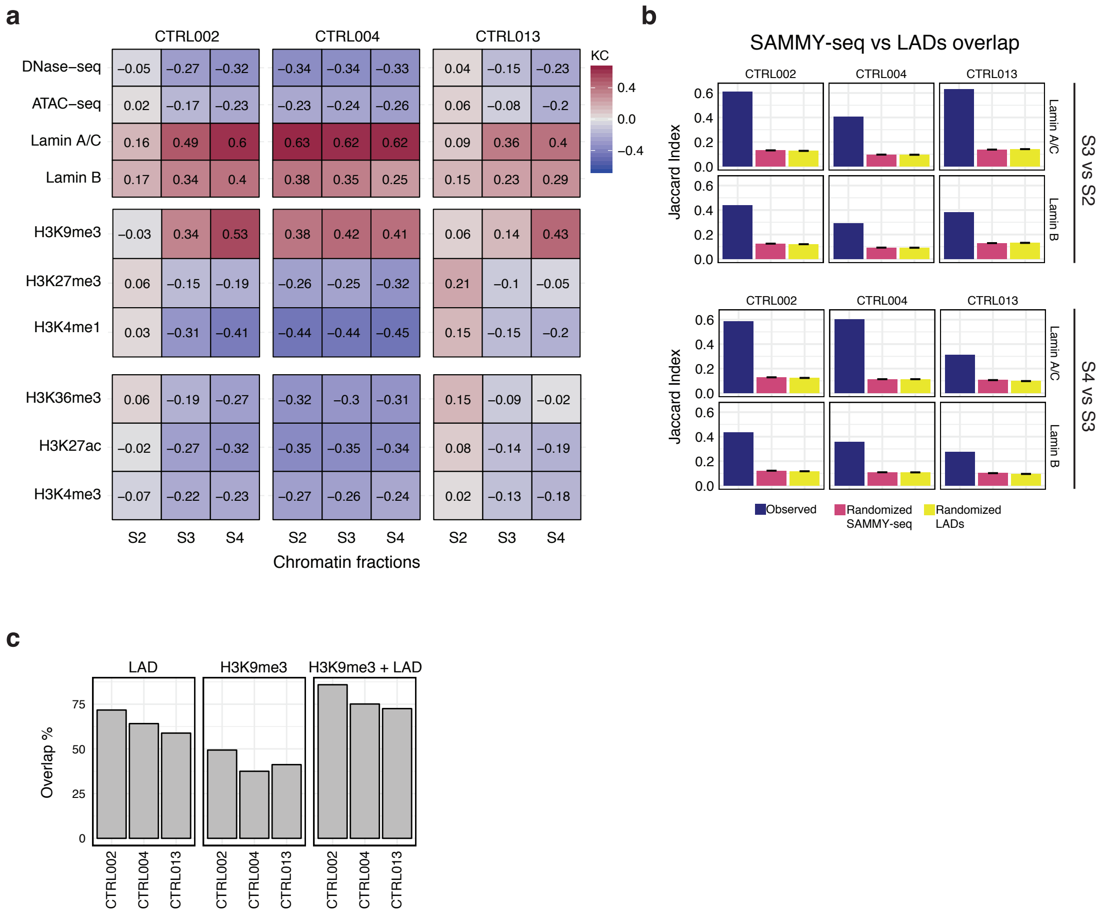

**Supplementary Figure 2 - Detailed characteristics and genome-wide association of SAMMY-seq fractions. a**, Genome-wide kernel correlation calculated by StereoGene, between read coverage in individual control samples chromatin fractions against ChIP-seq and other enrichment signals in ATAC-seq, DNase-seq, Lamin A/C and Lamin B; H3K27me3, H3K4me1 and H3K9me3; H3K36me3, H3K27ac and H3K4me3. In most samples the correlation is progressively increasing for closed chromatin marks or decreasing for open chromatin marks when considering fractions from S2 to S4. **b**, Overlap (Jaccard Index - JI) of control samples SAMMY-seq domains (S3 vs S2 or S4 vs S3) with LADs (Lamin A/C or Lamin B ChIP-seq enrichment domains). The observed JI (blue bar) is compared to median JI across 10,000 randomizations of SAMMY-seq domains (pink bar) or LADs (yellow bar) positions along the genome. All Jaccard Indexes are significantly higher than the randomized values (all empirical p-values < 0.0001). The black rectangle over the randomized values shows the +/- 2 standard error range. **c**, Percentage of LAD, H3K9me3 or LAD+H3K9me3 regions detected by the S4 vs S2 SAMMY-seq domains. LADs were defined based on[27], while H3K9me3 regions were originating from the Roadmap Epigenomics E055 sample[32].

### Supplementary Figure 3

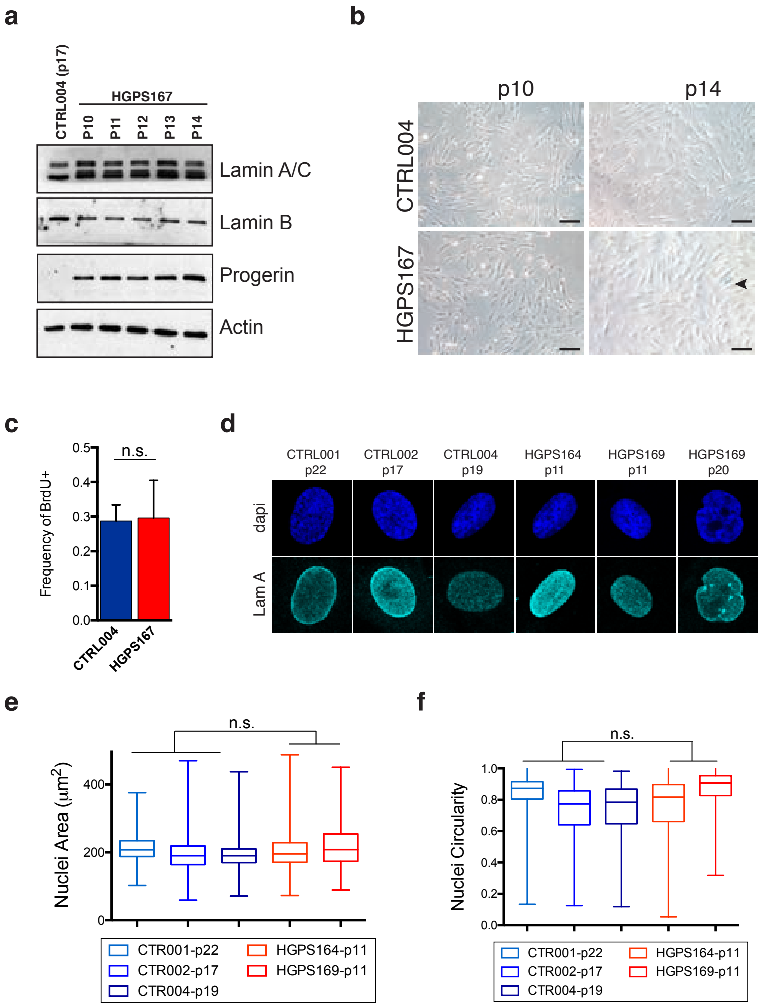

**Supplementary Figure 3 - Characteristics of control and progeria fibroblast cells. a**, Western blots for total extract from CTRL and HGPS fibroblasts at different cell culture passages (P10-14) hybridized with indicated antibodies. **b**, Representative fields of SA-β-gal activity in CTRL and HGPS fibroblasts at p10 and p14. Scale bar, 50 μm. **c**, Bar plot reporting the proliferation rate in control vs HGPS fibroblasts estimated as percentage of BrdU positive cells with respect to the total number of nuclei (n > 550). The bars report averages of two independent experiments, whiskers represent standard deviation. **d**, Representative images of Lamin A/DAPI immunofluorescence analysis on control or HGPS fibroblasts at various passages (indicated with p). **e**, **f**, Quantification of nuclei area (panel e) and circularity (panel f) of CTRL and HGPS fibroblasts. Number of analyzed nuclei: CTRL001: 660; CTRL002: 1253; CTRL004: 885; HGPS164: 990; HGPS169-p11: 312; HGPS169-p20: 150.
Comparisons were done using a two-tailed t-test in (c) and nested anova in (e, f). n.s.: not significant.

### Supplementary Figure 4

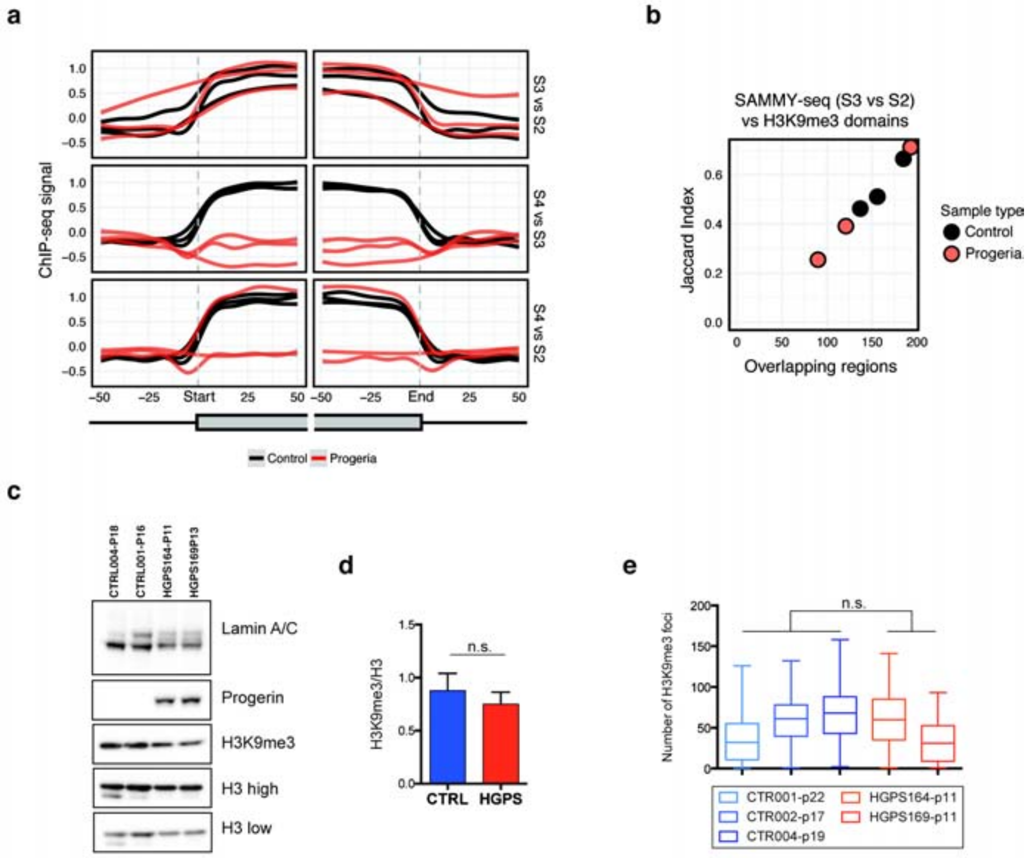

**Supplementary Figure 4 - Additional analysis on H3K9me3 patterns. a**, Average ChIP-seq enrichment signal for H3K9me3 is reported around the SAMMY-seq domain borders start (left side plots) or end (right side plots) for domains detected in each control (black lines) and progeria (red lines) sample. H3K9me3 ChIP-seq was obtained for each sample individually. Results for each set of SAMMY-seq enrichment domains are reported (S3 vs S2 top, S4 vs S3 middle, S4 vs S2 bottom) using a +/-50 bins window (10Kb bin size) centered on the start or end domain border positions (vertical dashed grey line). **b**, Overlap of H3K9me3 enriched domains and SAMMY-seq domains (S3 vs S2) for control (black dots) or HGPS (red dots) samples (JI on y-axis, number of overlapping regions on x-axis). **c**, Western blots for total extract from control or HGPS fibroblasts at early passages hybridized with indicated antibodies. Histone H3 was used as loading controls for H3K9me3 quantification. **d**, The graph shows quantifications of H3K9me3 levels normalized on H3. Data points were generated from an average of at least 10 biological replicates. **e**, Quantification of number per nucleus of H3K9me3 foci of CTRL and HGPS fibroblasts. Number of analyzed nuclei: CTRL001: 242; CTRL002: 649; CTRL004: 518; HGPS164: 550; HGPS169: 65.
Comparisons were done using a two-tailed t-test in (d) and nested anova in (e). n.s.: not significant.

### Supplementary Figure 5

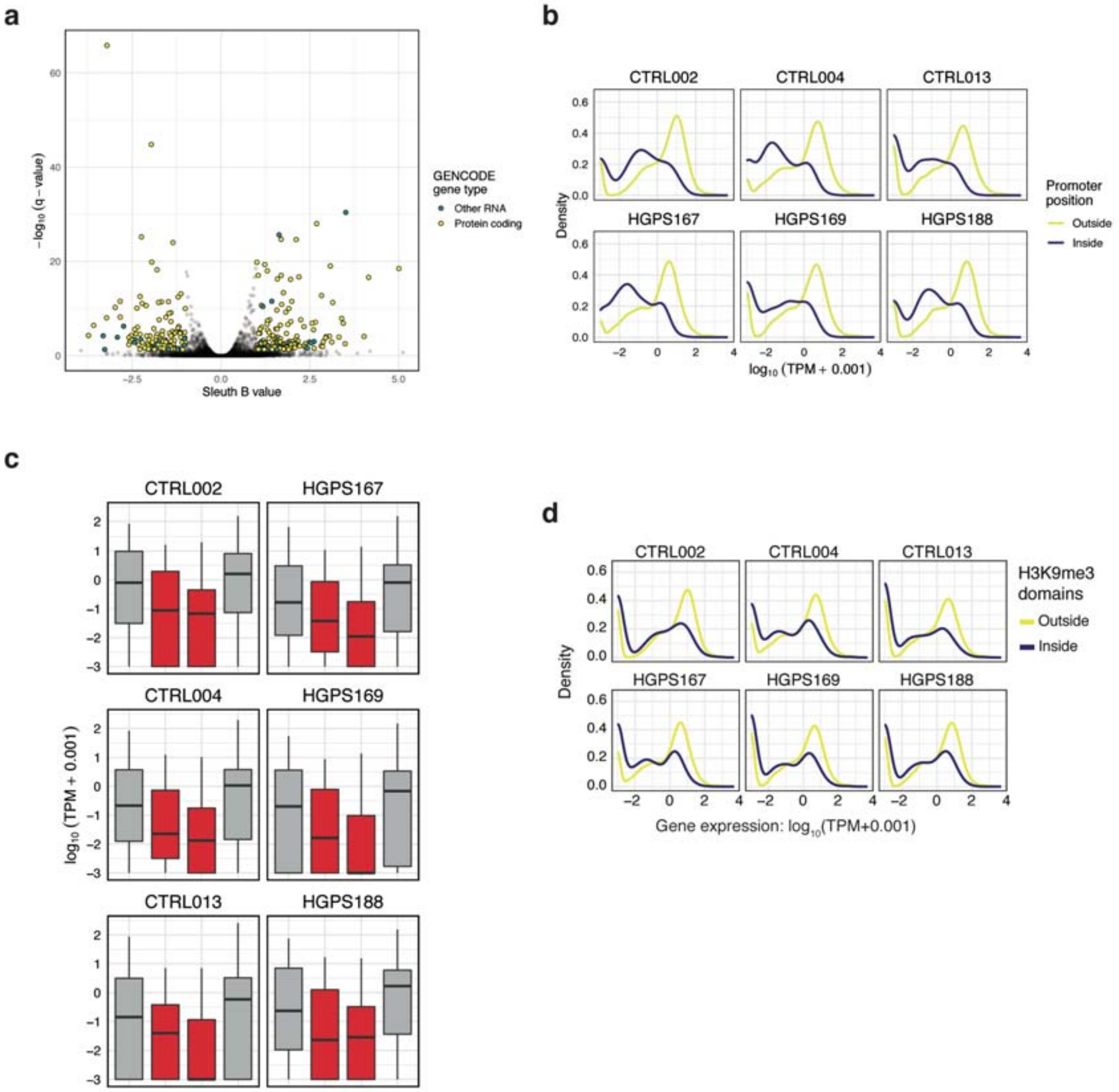

**Supplementary Figure 5 – Expression analysis in control and progeria samples. a**, Volcano plot of gene level differential expression. The x axis shows the sleuth b value, while the y axis shows $-\log_{10}$(q-value) of genes. Genes highlighted in green or yellow show significant changes (|b| > 1 and q-value < 0.05). The yellow color refers to protein coding genes based on GENCODE v27 annotation, green refers to all other types of non protein coding genes. **b**, Normalized gene expression distribution for protein coding genes separated based on their promoter being inside or outside of the consensus SAMMY-seq domains (S4 vs S2) of all controls. Genes with promoters outside the domains have higher expression (Wilcoxon rank sum test p-values < 0.01). **c**, Boxplot of normalized gene expression distribution for protein coding genes flanking consensus SAMMY-seq domain borders (S4 vs S2) of all controls. The gray boxplots show genes located in the outer 500Kb flanking regions (upstream of start or downstream of end), the red boxplots show genes in the inner 500Kb flanking regions (all Wilcoxon rank sum test p-values < 0.01). **d**, Gene expression distribution for protein coding genes separated based on their position being inside or outside of the H3K9me3 peaks in each sample. All samples show a significant difference in expression (Wilcoxon rank sum test p-values < 0.01).

### Supplementary Figure 6

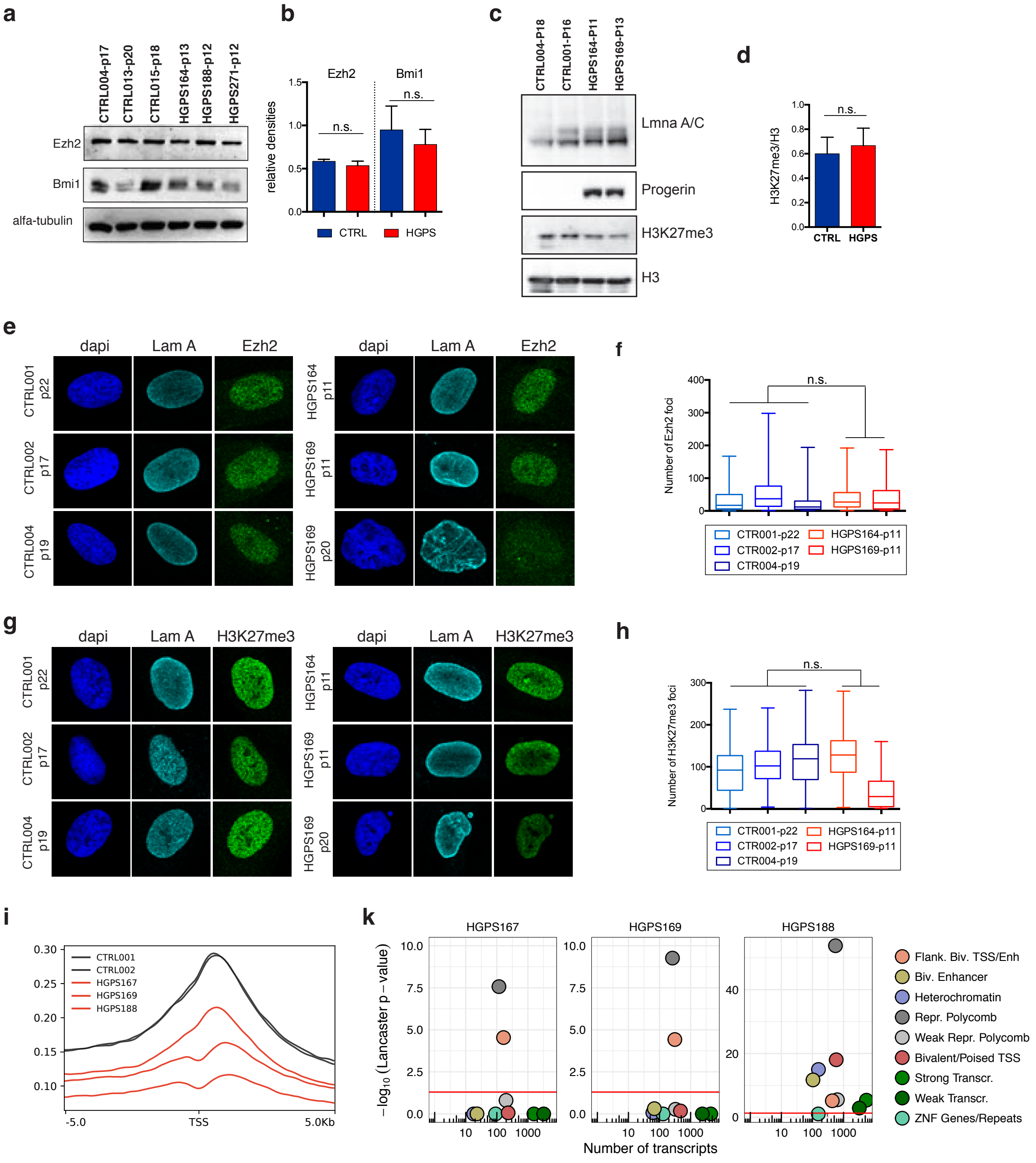

**Supplementary Figure 6 - Additional analysis on H3K27me3 patterns.** **a**, Western blots for total extract from control or HGPS fibroblasts at early passages hybridized with indicated antibodies. Alpha-tubulin was used as loading control. **b**, The graph shows the average of Ezh2 and Bmi1 levels in (a) normalized on Alpha-tubulin. **c**, Western blots for total extract from control or HGPS fibroblasts at early passages hybridized with indicated antibodies. Histone H3 was used as loading controls for H3K27me3 quantification. **d**, The graph shows quantifications of H3K27me3 levels normalized on H3. Data points were generated from an average of at least 4 biological replicates. **e**, Representative images of Ezh2/Lamin A immunofluorescence analysis on control or HGPS fibroblasts. **f** Quantification of number per nucleus of Ezh2 foci of CTRL and HGPS fibroblasts. Number of analyzed nuclei: CTRL001: 225; CTRL002: 265; CTRL004: 202; HGPS164: 188; HGPS169: 118. **g**, Representative images of H3K27me3/Lamin A immunofluorescence analysis on control or HGPS fibroblasts. **h**, Quantification of number per nucleus of H3K27me3 foci of CTRL and HGPS fibroblasts. Number of analyzed nuclei: CTRL001: 221; CTRL002: 339; CTRL004: 165; HGPS164: 251; HGPS169: 129. **i**, H3K27me3 signal distribution around the TSS region of all protein coding genes, based on GENCODE v27 annotation, calculated by deepTools using the genome-wide signal from SPP (see Online methods for details). The x-axis represents relative genomic position around the TSS (+/- 5Kb), and the y-axis represents average signal intensity. **k**, Transcripts differential expression (up-regulation) was assessed by comparing individual HGPS samples against the group of controls, then p-values were aggregated based on their chromatin state, considering only genes in SAMMY-seq domains (S4 vs S3) (see Online methods for details). The plot reports the number of transcripts (x-axis) and significance (Lancaster method aggregated p-values – y-axis) for each chromatin state (color legend) as defined by Roadmap Epigenomics[32].
Comparisons were done using a two-tailed t-test in (b, d) and nested anova in (f, h). n.s.: not significant.
